# Large-scale analysis of DNA methylation identifies cellular alterations in blood from psychosis patients and molecular biomarkers of treatment-resistant schizophrenia

**DOI:** 10.1101/2020.04.05.026211

**Authors:** Eilis Hannon, Emma L Dempster, Georgina Mansell, Joe Burrage, Nick Bass, Marc M Bohlken, Aiden Corvin, Charles J Curtis, David Dempster, Marta Di Forta, Timothy G Dinan, Gary Donohoe, Fiona Gaughran, Michael Gill, Amy Gillespie, Cerisse Gunasinghe, Hilleke E Hulshoff, Christina M Hultman, Viktoria Johansson, Rene S Kahn, Jaakko Kaprio, Gunter Kenis, Kaarina Kowalec, James MacCabe, Colm McDonald, Andew McQuillin, Derek W Morris, Kieran C Murphy, Collette Mustard, Igor Nenadic, Michael C O’Donovan, Diego Quattrone, Alexander L Richards, Bart PF Rutten, David St Clair, Sebastian Therman, Timothea Toulopoulou, Jim Van Os, John L Waddington, Wellcome Trust Case Control Consortium 2, CREeTable AR consortium, Patrick Sullivan, Evangelos Vassos, Gerome Breen, David Andrew Collier, Robin Murray, Leonard S Schalkwyk, Jonathan Mill

## Abstract

**Objective:** Psychosis - a complex and heterogeneous neuropsychiatric condition characterized by hallucinations and delusions - is a common feature of schizophrenia. There is evidence for altered DNA methylation (DNAm) associated with schizophrenia in both brain and peripheral tissues. We aimed to undertake a systematic analysis of variable DNAm associated with psychosis, schizophrenia, and treatment-resistant schizophrenia, also exploring measures of biological ageing, smoking, and blood cell composition derived from DNAm data to identify molecular biomarkers of disease.

**Methods:** We quantified DNAm across the genome in blood samples from 4,483 participants from seven case-control cohorts including patients with schizophrenia or first-episode psychosis. Measures of biological age, cellular composition and smoking status were derived from DNAm data using established algorithms. DNAm and derived measures were analyzed within each cohort and the results combined by meta-analysis.

**Results:** Psychosis cases were characterized by significant differences in measures of blood cell proportions and elevated smoking exposure derived from the DNAm data, with the largest differences seen in treatment-resistant schizophrenia patients. DNAm at 95 CpG sites was significantly different between psychosis cases and controls, with 1,048 differentially methylated positions (DMPs) identified between schizophrenia cases and controls. Schizophrenia-associated DMPs colocalize to regions identified in genetic association studies, with genes annotated to these sites enriched for pathways relevant to disease. Finally, a number of the schizophrenia associated differences were only present in the treatment-resistant schizophrenia subgroup.

**Conclusions:** We show that DNAm data can be leveraged to derive measures of blood cell counts and smoking that are strongly associated with psychosis. Our DNAm meta-analysis identified multiple DMPs associated with both psychosis and a more refined diagnosis of schizophrenia, with evidence for differential methylation associated with treatment-resistant schizophrenia that potentially reflects exposure to clozapine.

## Introduction

Psychosis is a complex and heterogeneous neuropsychiatric condition, characterized by hallucinations and delusions. Episodic psychosis and altered cognitive function are major features of schizophrenia, a severe neurodevelopmental disorder that contributes significantly to the global burden of disease ^1^. Schizophrenia is highly heritable ^2,3^ and recent genetic studies have indicated a complex polygenic architecture involving hundreds of genetic variants that individually confer a minimal increase on the overall risk of developing the disorder^4^. Large-scale genome-wide association studies (GWAS) have identified approximately 160 regions of the genome harboring common variants robustly associated with the diagnosis of schizophrenia, with evidence for a substantial polygenic component in signals that individually fall below genome-wide levels of significance ^5,6^. As the majority of schizophrenia-associated variants do not directly index coding changes affecting protein structure, there remains uncertainty about the causal genes involved in disease pathogenesis, and how their function is dysregulated ^7^.

A major hypothesis is that GWAS variants predominantly act to influence the regulation of gene expression. This hypothesis is supported by an enrichment of schizophrenia associated variants in core regulatory domains (e.g. active promotors and enhancers)^8^. As a consequence, there has been growing interest in the role of epigenetic variation in the molecular etiology of schizophrenia. DNA methylation is the best-characterized epigenetic modification, acting to influence gene expression via disruption of transcription factor binding and recruitment of methyl-binding proteins that initiate chromatin compaction and gene silencing. Despite being traditionally regarded as a mechanism of transcriptional repression, DNA methylation is actually associated with both increased and decreased gene expression^9^, and other genomic functions including alternative splicing and promoter usage^10^. We previously demonstrated how DNA methylation is under local genetic control^11,12^, identifying an enrichment of DNA methylation quantitative trait loci (mQTL) among genomic regions associated with schizophrenia^11^. Furthermore, we have used mQTL associations to identify discrete sites of regulatory variation associated with schizophrenia risk variants implicating specific genes within these regions ^11-14^. Of note, epigenetic variation induced by non-genetic exposures has been hypothesized as another mechanism by which environmental factors can affect risk for neuropsychiatric disorders including schizophrenia^15^.

The development of standardized assays for quantifying DNA methylation at specific sites across the genome has enabled the systematic analysis of associations between methylomic variation and environmental exposures or disease^16^. Because DNA methylation is a dynamic process, these epigenome-wide association studies (EWAS) are more complex to design and interpret than GWAS^17-19^. As for observational epidemiological studies of exposures and outcomes, a number of potentially important confounding factors (e.g. tissue- or cell-type, age, sex, lifestyle exposures, medication, and disorder-associated exposures) that can directly influence DNA methylation need to be considered along with the possibility of reverse causation. Despite these difficulties, recent studies have identified schizophrenia-associated DNA methylation differences in analyses of post-mortem brain tissue^20-23^, and also detected disease-associated variation in peripheral blood samples from both schizophrenia-discordant monozygotic twin pairs ^24^ and clinically-ascertained case-control cohorts ^13,25,26^. We previously reported an EWAS of variable DNA methylation associated with schizophrenia in >1,700 individuals, meta-analyzing data from three independent cohorts and identifying methylomic biomarkers of disease^13^.Together these data support a role for differential DNA methylation in the molecular etiology of schizophrenia, although it is not clear whether disease-associated methylation differences are themselves secondary to the disorder itself, or a result of other schizophrenia-associated factors.

In this study we extend our previous analysis, quantifying DNA methylation across the genome in a total of 4,483 participants from seven independent case-control cohorts including patients with schizophrenia or first-episode psychosis (FEP). In each cohort, genomic DNA was isolated from whole blood and DNA methylation was quantified across the genome using either the Illumina Infinium HumanMethylation450 microarray (“450K array”) or the HumanMethylationEPIC microarray (“EPIC array”) (see **Methods**). We implemented a stringent pipeline to meta-analyze EWAS results across datasets to identify associations between psychosis cases and variation in DNA methylation. We show how DNA methylation data can be leveraged to identify biological (e.g. differential cell counts) and environmental (e.g. smoking) factors associated with psychosis, and present evidence for molecular variation associated with clozapine exposure in patients with treatment-resistant schizophrenia.

## Methods

### Cohort descriptions

#### University College London (UCL) samples

447 schizophrenia cases and 456 controls from the University College London schizophrenia sample cohort were selected for DNA methylation profiling. A full description of this cohort can be found elsewhere^27^ but briefly comprises of unrelated ancestrally matched cases and controls from the United Kingdom. Case participants were recruited from UK NHS mental health services with a clinical ICD-10 diagnosis of schizophrenia. All case participants were interviewed with the Schedule for Affective Disorders and Schizophrenia-Lifetime Version (SADS-L)^28^ to confirm Research Diagnostic Criteria (RDC) diagnosis. A control sample screened for an absence of mental health problems was recruited. Each control subject was interviewed to confirm that they did not have a personal history of an RDC defined mental disorder or a family history of schizophrenia, bipolar disorder, or alcohol dependence. UK National Health Service multicentre and local research ethics approval was obtained and all subjects signed an approved consent form after reading an information sheet.

#### Aberdeen samples

482 schizophrenia cases and 468 controls from the Aberdeen schizophrenia sample were selected for DNA methylation profiling. The Aberdeen case-control sample has been fully described elsewhere ^29^ but briefly contains schizophrenia cases and controls who have self-identified as born in the British Isles (95% in Scotland). All cases met the Diagnostic and Statistical Manual for Mental Disorders-IV edition (DSM-IV) and International Classification of Diseases 10th edition (ICD-10) criteria for schizophrenia. Diagnosis was made by Operational Criteria Checklist (OPCRIT). Controls were volunteers recruited through general practices in Scotland. Practice lists were screened for potentially suitable volunteers by age and sex and by exclusion of subjects with major mental illness or use of neuroleptic medication. Volunteers who replied to a written invitation were interviewed using a short questionnaire to exclude major mental illness in individual themselves and first-degree relatives. All cases and controls gave informed consent. The study was approved by both local and multiregional academic ethical committees.

#### Monozygotic twins discordant for schizophrenia

The monozygotic twin cohort is a multi-centre collaborative project aimed at identifying DNA methylation differences in monozygotic-twin pairs discordant for a diagnosis of schizophrenia. 96 informative twin-pairs (n = 192 individuals) were identified from European twin studies based in Utrecht (The Netherlands), Helsinki (Finland), London (United Kingdom), Stockholm (Sweden), and Jena (Germany). Of the monozygotic twin pairs utilized in the analysis, 75 were discordant for diagnosed schizophrenia, 6 were concordant for schizophrenia and 15 twin pairs were free of any psychiatric disease. Each twin study has been approved; ethical permission was given by the relevant local ethics committee and the participating twins have provided written informed consent.

#### Dublin samples

361 schizophrenia cases and 346 controls were selected from the Irish Schizophrenia Genomics consortium, a detailed description of this cohort can be found in the Morris et al manuscript ^30^. Briefly, participants, from the Republic of Ireland or Northern Ireland, were interviewed using a structured clinical interview and diagnosis of schizophrenia or a related disorder [schizoaffective disorder; schizophreniform disorder] was made by the consensus lifetime best estimate method using DSM-IV criteria. Control subjects were ascertained with written informed consent from the Irish GeneBank and represented blood donors from the Irish Blood Transfusion Service. Ethics Committee approval for the study was obtained from all participating hospitals and centres.

#### IoPPN samples

The IoPPN cohort comprises of 290 schizophrenia cases, 308 first episode psychosis (FEP) patients and 203 non-psychiatric controls recruited from the same geographical area into three studies via the South London & Maudsley Mental Health National Health Service (NHS) Foundation Trust. Established schizophrenia cases were recruited to the Improving Physical Health and Reducing Substance Use in Severe Mental Illness (IMPACT) study from three English mental health NHS services ^31^. First episode psychosis patients were recruited to the GAP study^32^ via in-patient and early intervention in psychosis community mental health teams. All patients aged 18–65 years who presented with a first episode of psychosis to the Lambeth, Southwark and Croydon adult in-patient units of the South London & Maudsley Mental Health NHS Foundation Trust between May 1, 2005, and May 31, 2011 who met ICD–10 criteria for a diagnosis of psychosis (codes F20–F29 and F30–F33). Clinical diagnosis was validated by administering the Schedules for Clinical Assessment in Neuropsychiatry (SCAN). Cases with a diagnosis of organic psychosis were excluded. Healthy controls were recruited into the GAP study from the local population living in the area served by the South London & Maudsley Mental Health NHS Foundation Trust, by means of internet and newspaper advertisements, and distribution of leaflets at train stations, shops and job centres. Those who agreed to participate were administered the Psychosis Screening Questionnaire^33^ and excluded if they met criteria for a psychotic disorder or reported to have received a previous diagnosis of psychotic illness. All participants were included in the study only after giving written, confirmed consent. The study protocol and ethical permission was granted by the Joint South London and Maudsley and the Institute of Psychiatry NHS Research Ethics Committee (17/NI/0011).

#### Sweden

190 schizophrenia cases and 190 controls from the Sweden Schizophrenia Study (S3) [31] were selected for DNA methylation profiling details of which have been described previously [2]. Briefly, S3 is a population-based cohort of individuals born in Sweden including 4,936 SCZ cases and 6,321 healthy controls recruited between 2004 and 2010. SCZ cases were identified from the Sweden Hospital Discharge Register [32, 33] with ≥2 hospitalizations with an ICD discharge diagnosis of SCZ or schizoaffective disorder (SAD) [34]. This operational definition of SCZ was validated in clinical, epidemiological, genetic epidemiological, and genetic studies [31]. More generally, the Hospital Discharge Register has high agreement with medical [32, 33] and psychiatric diagnoses [35]. Controls were also selected through Swedish Registers and were group-matched by age, sex and county of residence and had no lifetime diagnoses of SCZ, SAD, or bipolar disorder or antipsychotic prescriptions. Blood samples were drawn at enrolment. All subjects were 18 years of age or older and provided written informed consent. Ethical permission was obtained from the Karolinska Institutet Ethical Review Committee in Stockholm, Sweden.

#### The European Network of National Schizophrenia Networks Studying Gene-Environment Interactions (EU-GEI) cohort

458 first-episode psychosis (FEP) cases and 558 controls from the incidence and case-control work package (WP2) of the European Network of National Schizophrenia Networks Studying Gene-Environment Interactions (EU-GEI) cohort were selected for DNA methylation profiling ^34^. Patients presenting with FEP were identified, between 1/5/2010 and 1/4/2015, by trained researchers who carried out regular checks across the 17 catchment area Mental Health Services across 6 European countries. FEP were included if a) age 18-64 years and b) resident within the study catchment areas at the time of their first presentation, and with a diagnosis of psychosis (ICD-10 F20-33). Using the Operational Criteria Checklist algorithm ^35,36,35,36^(35, 36) all cases interviewed received a research-based diagnosis. FEPs were excluded if a) previously treated for psychosis, b) they met criteria for organic psychosis (ICD-10: F09), or for a diagnosis of transient psychotic symptoms resulting from acute intoxication (ICD-10: F1X.5). FEP were approached via their clinical team and invited to participate in the assessment. Random and Quota sampling strategies were adopted to guide the recruitment of controls from each of the sites. The most accurate local demographic data available were used to set quotas for controls to ensure the samples’ representativeness of each catchment area’s population at risk. Controls were excluded if a) they had received a diagnosis of, b) and/or treatment for, psychotic disorder. All participants provided informed, written consent. Ethical approval was provided by relevant research ethics committees in each of the study sites.

#### Genome-wide quantification of DNA methylation

Approximately 500ng of blood-derived DNA from each sample was treated with sodium bisulfite in duplicate, using the EZ-96 DNA methylation kit (Zymo Research, CA, USA). DNA methylation was quantified using either the Illumina Infinium HumanMethylation450 BeadChip (Illumina Inc, CA, USA) or Illumina Infinium HumanMethylationEPIC BeadChip (Illumina Inc, CA, USA) run on an Illumina iScan System (Illumina, CA, USA) using the manufacturers’ standard protocol. Samples were batched by cohort and randomly assigned to chips and plates to ensure equal distribution of cases and controls across arrays and minimize batch effects. For the monozygotic Twin cohort, both members of the same twin pair were run on the same chip. A fully methylated control sample (CpG Methylated HeLa Genomic DNA; New England BioLabs, MA, USA) was included in a random position on each plate to facilitate plate tracking. Signal intensities were imported in R programming environment using the *methylumIDAT* function in the *methylumi* package ^37^. Our stringent quality control pipeline included the following steps: 1) checking methylated and unmethylated signal intensities, excluding samples where this was < 2500; 2) using the control probes to ensure the sodium bisulfite conversion was successful, excluding any samples with median < 90; 3) identifying the fully methylated control sample was in the correct location; 4) all tissues predicted as of blood origin using the tissue prediction from the Epigenetic Clock software (https://DNAmAge.genetics.ucla.edu/) ^38^; 5) multidimensional scaling of sites on X and Y chromosomes separately to confirm reported gender; 6) comparison with genotype data across SNP probes; 7) *pfilter* function from wateRmelon package ^39^ to exclude samples with > 1% of probes with detection *P*-value > 0.05 and probes with > 1% of samples with detection *P*-value > 0.05. PCs were used (calculated across all probes) to identify outliers, samples > 2 standard deviations from the mean for both PC1 and PC2 were removed. An additional QC step was performed in the Twins cohort using the 65 SNP probes to confirm that twins were genetically identical. Normalization of the DNA methylation data was performed used the *dasen* function in the *wateRmelon* package^39^. As cell count data were not available for these DNA samples these were estimated from the 450K DNA methylation data using both the Epigenetic Clock software ^38^ and Houseman algorithm ^40,41^, including the 7 variables recommended in the documentation for the Epigenetic Clock in the regression analysis. For cohorts with the EPIC array DNA methylation data, we were only able to generate the 6 cellular composition variables using the Houseman algorithm^40,41^, which were included as covariates.

Similarly as smoking data was incomplete for the majority of cohorts, we calculated a smoking score from the data using the method described by Elliot et al^42^ and successfully used in our previous (Phase 1) analyses^13^. Raw and processed data for the UCL, Aberdeen and Dublin cohorts are available through GEO accession numbers GSE84727, GSE80417, and GSE147221 respectively.

#### Data analysis

All analyses were performed with the statistical language R unless otherwise stated. Custom code for all steps of the analysis are available on GitHub: (https://github.com/ejh243/SCZEWAS/tree/master/Phase2).

#### Comparison of derived estimates of cellular composition and tobacco smoking

A linear regression model was used to test for differences in ten cellular composition variables estimated from the DNA methylation data, reflecting either proportion or abundance of blood cell types. These estimated cellular composition variables were regressed against case/control status with covariates for age, sex and smoking. Estimated effects and standard errors were combined across the cohorts using a random effect meta-analysis implemented with the meta package^43^. The same methodology was used to test for differences in the DNA methylation derived smoking score between cases and controls including covariates for age and sex. P values are from two-sided tests.

#### Within-cohort EWAS analysis

A linear regression model was used to test for differentially methylated sites associated with schizophrenia or first episode psychosis. DNA methylation values for each probe were regressed against case/control status with covariates for age, sex, cell composition, smoking status and batch. For the EU-GEI cohort there was an additional covariate for contributing study. For the Twins cohort, a linear model was used to generate regression coefficients, but *P*-values were calculated with clustered standards errors using the *plm* package ^44^, recognising individuals from the same twin pair.

#### Within-patient EWAS of clozapine prescription

Four individual cohorts (UCL, Aberdeen, IoPPN and Sweden) had information on medication and/or clozapine exposure and were included in the treatment-resistant schizophrenia (TRS) EWAS. TRS patients were defined as any case that had ever been prescribed clozapine, and non-TRS patients were defined as schizophrenia cases that had no record of being prescribed clozapine. Within each cohort DNA methylation values for each probe were regressed against TRS status with covariates for age, sex, cell composition, smoking status, and batch as described for the case control EWAS.

#### Meta-analysis

The EWAS results from each cohort were processed using the *bacon* R package^45^, which uses a Bayesian method to adjust for inflation in EWAS P-values. All probes analysed in at least two studies were taken forward for meta-analysis. This was performed using the *metagen* function in the R package meta^43^, using the effect sizes and standard errors adjusted for inflation from each individual cohort to calculate weighted pooled estimates and test for significance. P-values are from two-sided tests and significant DMPs were identified from a random effects model at a significance threshold of 9×10^−8^, which controls for the number of independent tests performed when analysis data generated with the EPIC array^46^. DNA methylation sites were annotated with location information for genome build hg19 using the Illumina manifest files (CHR and MAPINFO).

#### Overlap with schizophrenia GWAS loci

The GWAS regions were taken from the largest published schizophrenia GWAS to date by Pãrdinas et al.^6^ made available through the Psychiatric Genomics Consortium (PGC) website (https://www.med.unc.edu/pgc/results-and-downloads). Briefly, regions were defined by performing a “clumping” procedure on the GWAS *P*-values to collapse multiple correlated signals (due to linkage disequilibrium) surrounding the index SNP (i.e. with the smallest P-value) into a single associated region. To define physically distinct loci, those within 250kb of each other were subsequently merged to obtain the final set of GWAS regions. The outermost SNPs of each associated region defined the start and stop parameters of the region. Using the set of 158 schizophrenia-associated genomic loci we used Brown’s method ^47^ to calculate a combined P-value across all probes located within each region (based on hg19) using the probe-level P-values and correlation coefficients between all pairs of probes calculated from the DNA methylation values. As correlations between probes could only be calculated using probes profiled on the same array, this analysis was limited to probes included on the EPIC array. Correlations between probes were calculated within the EU-GEI cohort as this had the largest number of samples.

#### Enrichment analyses

Enrichment of the heritability statistics of DMPs was performed against a background set of probes selected to match the distribution of the test set for both mean and standard deviation. This was achieved by splitting all probes into 10 equally sized bins based on their mean methylation level and ten equally sized bins based on their standard deviation, to create a matrix of 100 bins. After counting the number of DMPs within each bin, we selected the same number of probes from each bin for the background comparison set. This was repeated multiple times, without replacement, until all the probes from at least one bin were selected giving the maximum possible number of background probes (n = 42,968) such that they matched the characteristics of the test set of DMPs.

#### Gene ontology analysis

Illumina UCSC gene annotation, which is derived from the genomic overlap of probes with RefSeq genes or up to 1500bp of the transcription start site of a gene, was used to create a test gene list from the DMPs for pathway analysis. Where probes were not annotated to any gene (i.e. in the case of intergenic locations) they were omitted from this analysis, and where probes were annotated to multiple genes, all were included. A logistic regression approach was used to test if genes in this list predicted pathway membership, while controlling for the number of probes that passed quality control (i.e. were tested) annotated to each gene. Pathways were downloaded from the GO website (http://geneontology.org/) and mapped to genes including all parent ontology terms. All genes with at least one 450K probe annotated and mapped to at least one GO pathway were considered. Pathways were filtered to those containing between 10 and 2000 genes. After applying this method to all pathways, the list of significant pathways (P < 0.05) was refined by grouping to control for the effect of overlapping genes. This was achieved by taking the most significant pathway, and retesting all remaining significant pathways while controlling additionally for the best term. If the test genes no longer predicted the pathway, the term was said to be explained by the more significant pathway, and hence these pathways were grouped together. This algorithm was repeated, taking the next most significant term, until all pathways were considered as the most significant or found to be explained by a more significant term.

## RESULTS

### Study overview and cohort characteristics

We quantified DNA methylation in samples derived from peripheral venous whole blood in seven independent psychosis case-control cohorts (total n = 4,483; 2,379 cases and 2,104 controls). These cohorts represent a range of study designs and recruitment strategies and were initially designed to explore different clinical and etiological aspects of schizophrenia (see **Methods** and **Table 1**); they include studies of first episode psychosis (EU-GEI and IoPPN), established schizophrenia and/or clozapine usage (UCL, Aberdeen, Dublin, IoPPN), mortality in schizophrenia (Sweden), and a study of twins from monozygotic pairs discordant for schizophrenia (Twins). All cohorts were characterised by a higher proportion of male participants (range = 52.1–71.1% male, pooled mean = 62.6% male, **Table 1**) than females. Although there was an overall significantly higher proportion of males amongst cases compared to controls (P = 9.35×10^−10^), consistent with prevalence rates ^48,49^, there was significant heterogeneity in the sex by diagnosis proportions across different cohorts (P = 4.01×10^−61^) with the overall excess of male patients driven by two cohorts (UCL (P = 3.81×10^−13^) and EU-GEI (P = 3.68×10^−7^)). Most cohorts were enriched for young and middle-aged adults although there was considerable heterogeneity across the studies reflecting the differing sampling strategies (**Table 1**). For example, the IoPPN cohort has the lowest average age, reflecting the inclusion of a large number of first episode psychosis (FEP) patients (mean = 34.9 years; SD = 12.42 years)^50^. In contrast, individuals in the Sweden cohort were older (mean = 60.0 years; SD = 8.9 years)^51^. There was no overall difference in mean age between cases and controls (**eFigure 1**; P = 0.975), although differences were apparent in individual cohorts; in UCL (mean difference = 6.8 years; P = 6.55×10^−9^) and IoPPN (mean difference = 6.2 years; P = 1.46×10^−11^) patients were significantly older than controls, while in the EU-GEI (mean difference = -7.9 years; P = 1.24×10^−22^) and the Sweden cohort (mean difference = -7.3 years; P = 1.05×10^−16^) the cases were significantly younger.

**Table 1.** Summary of cohort demographics included in schizophrenia EWAS. FEP – first episode psychosis.

| **Cohort** |  | **UCL** | **Aberdeen** | **Twins** | **IoPPN** | **Dublin** | **EUGEI** | **Sweden** | **Combined** |
| --- | --- | --- | --- | --- | --- | --- | --- | --- | --- |
| Total sample |  | 675 | 847 | 192 | 800 | 679 | 912 | 378 | 4483 |
| % cases |  | 52.3 | 48.9 | 45.3 | 74.6 | 51.3 | 42.9 | 50.0 | 53.1 |
| % schizophrenia |  | 52.3 | 48.9 | 45.3 | 36.3 | 51.3 | 0.0 | 50.0 | 37.5 |
| % first episode psychosis |  | 0.0 | 0.0 | 0.0 | 38.4 | 0.0 | 42.9 | 0.0 | 15.6 |
| % Males | All | 58.7 | 71.1 | 52.1 | 63.0 | 71.0 | 54.4 | 59.5 | 62.6 |
|  | Cases | 72.0 | 68.4 | 54.0 | 65.3 | 71.6 | 64.2 | 60.3 | 66.8 |
|  | Controls | 44.1 | 73.7 | 50.5 | 56.2 | 70.4 | 47.0 | 58.7 | 57.8 |
|  | Chi-square test<br>P value | 3.81E-13 | 0.103 | 0.730 | 0.024 | 0.804 | 3.68E-07 | 0.834 | 9.35E-10 |
| Age<br>(years) | Mean | 40.4 | 44.6 | 35.3 | 28.8 | 41.7 | 35.3 | 60.0 | 40.5 |
|  | SD | 15.0 | 12.9 | 10.8 | 9.46 | 12.0 | 12.8 | 8.86 | 14.7 |
|  | Mean in<br>controls | 43.7 | 44.2 | 37.9 | 27.8 | 41.4 | 30.7 | 56.3 | 41.6 |
|  | Mean in cases | 36.8 | 44.9 | 33.3 | 30.3 | 42.0 | 38.7 | 63.7 | 39.4 |
|  | T-test Pvalue | 6.55E-09 | 0.529 | 0.033 | 0.007 | 0.505 | 1.24E-22 | 1.05E-16 |  |

**Figure 1.**
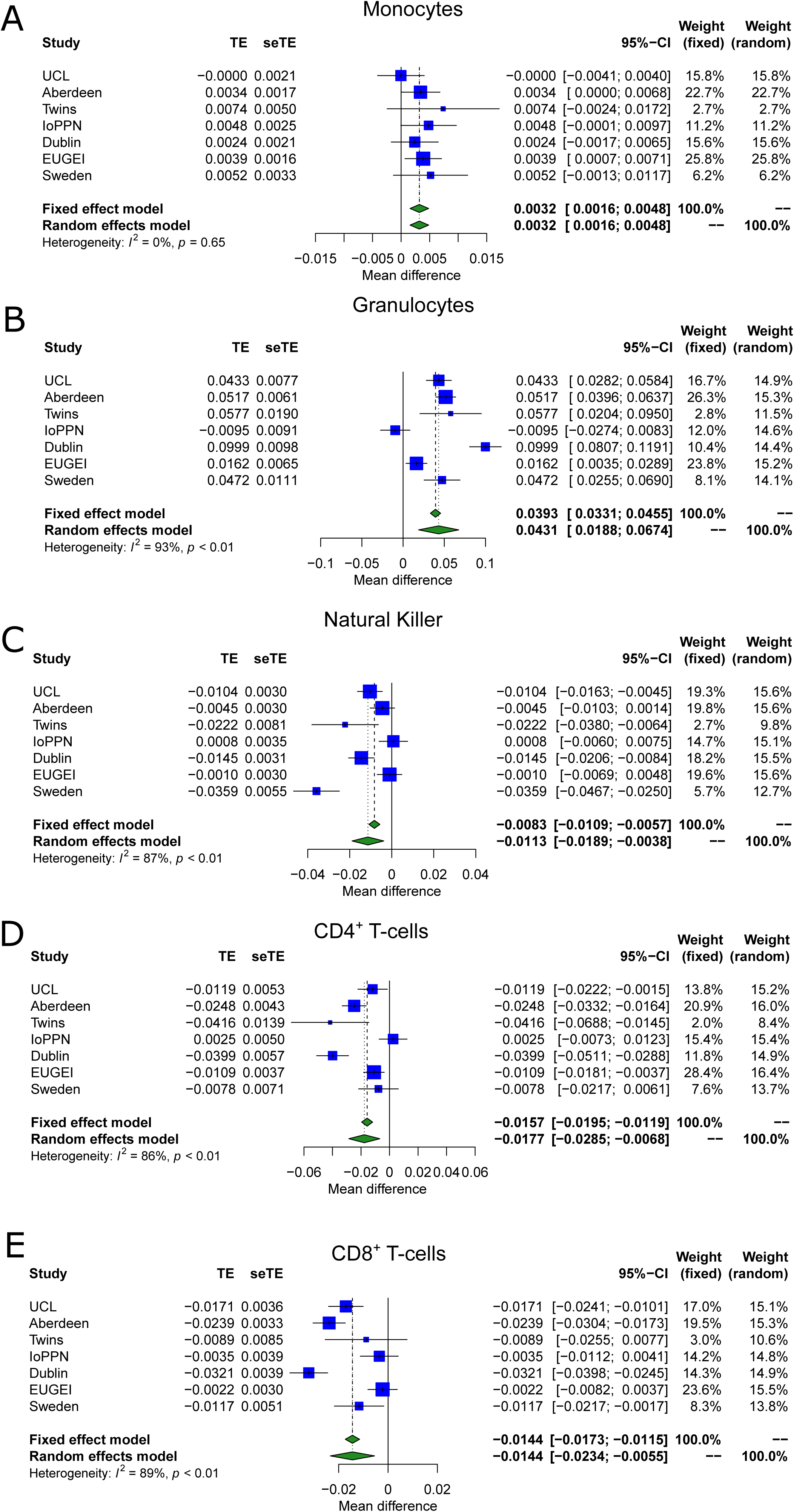
DNA methylation data highlight that schizophrenia cases are characterized by altered blood cell proportions. Shown are forest plots from meta-analyses of differences in blood cell proportions derived from DNA methylation data between psychosis patients and controls for **A**) monocytes **B)** granulocytes **C)** natural killer cells **D**) CD4+ T-cells and **E**) CD8+ T-cells. TE – treatment effect i.e. the mean difference between cases and controls, seTE – standard error of the treatment effect.

### Psychosis patients are characterized by differential blood cell proportions and smoking levels using measures derived from DNA methylation data

A number of robust statistical classifiers have been developed to derive estimates of both biological phenotypes (e.g. age ^38,52,53^ and the proportion of different blood cell types in a whole blood sample ^40,41^) and environmental exposures (e.g. tobacco smoking ^42,54^) from DNA methylation data. These estimates can be used to identify differences between groups and are often included as covariates in EWAS analyses where empirically-measured data is not available. For each individual included in this study we calculated two measures of “epigenetic age” from the DNA methylation data; DNAmAge using the Horvath multi-tissue clock, which was developed to predict chronological age ^38^, and PhenoAge, which was developed as biomarker of advanced biological aging ^55^. We found a strong correlation between reported age and both derived age estimates across the cohorts (Pearson correlation coefficient range 0.821-0.928 for DNAmAge (**eFigure 2**) and 0.795-0.910 for PhenoAge (**eFigure 3**)) and no evidence for age acceleration - i.e. the difference between epigenetic age and chronological age - between patients with psychosis and controls ^51^ (**eFigure 2** and **eFigure 3**).

**Figure 2.**
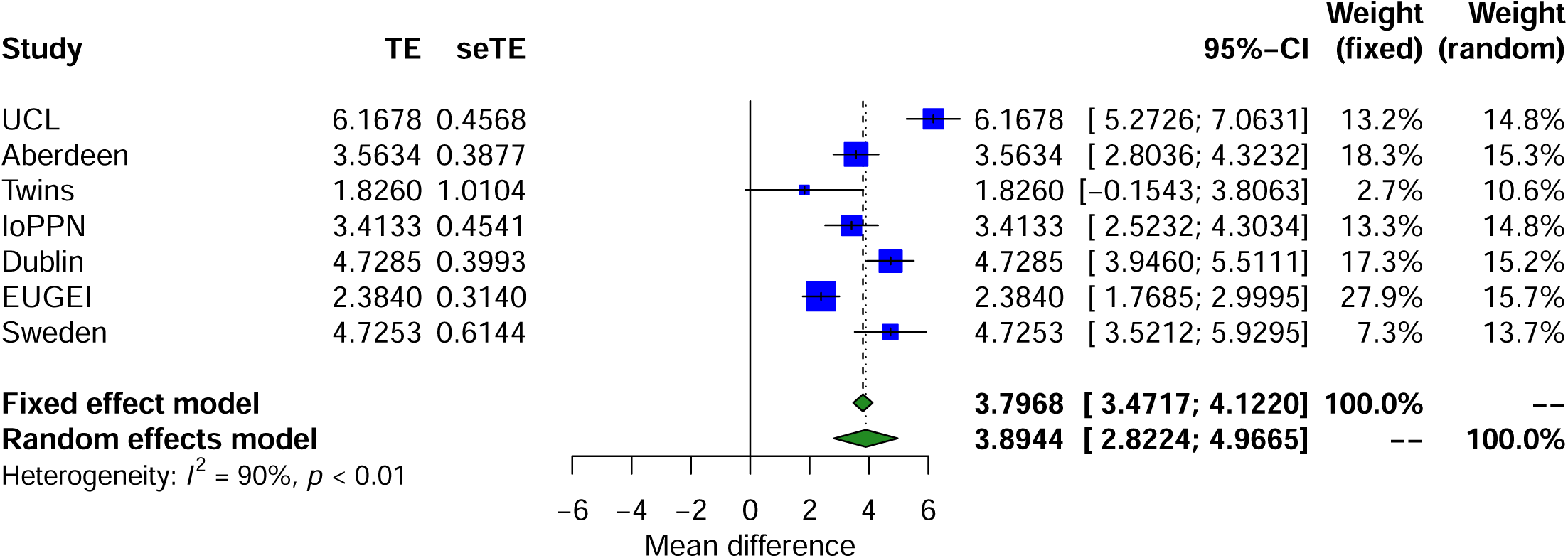
DNA methylation data highlight that psychosis patients are characterized by an elevated exposure to tobacco smoking. Forest plot from a meta-analysis of differences in smoking score derived from DNA methylation data between psychosis patients and controls. The smoking score was calculated from DNA methylation data using the method described by Elliott et al^42^. TE – treatment effect i.e. the mean difference between cases and controls, seTE – standard error of the treatment effect.

**Figure 3.**
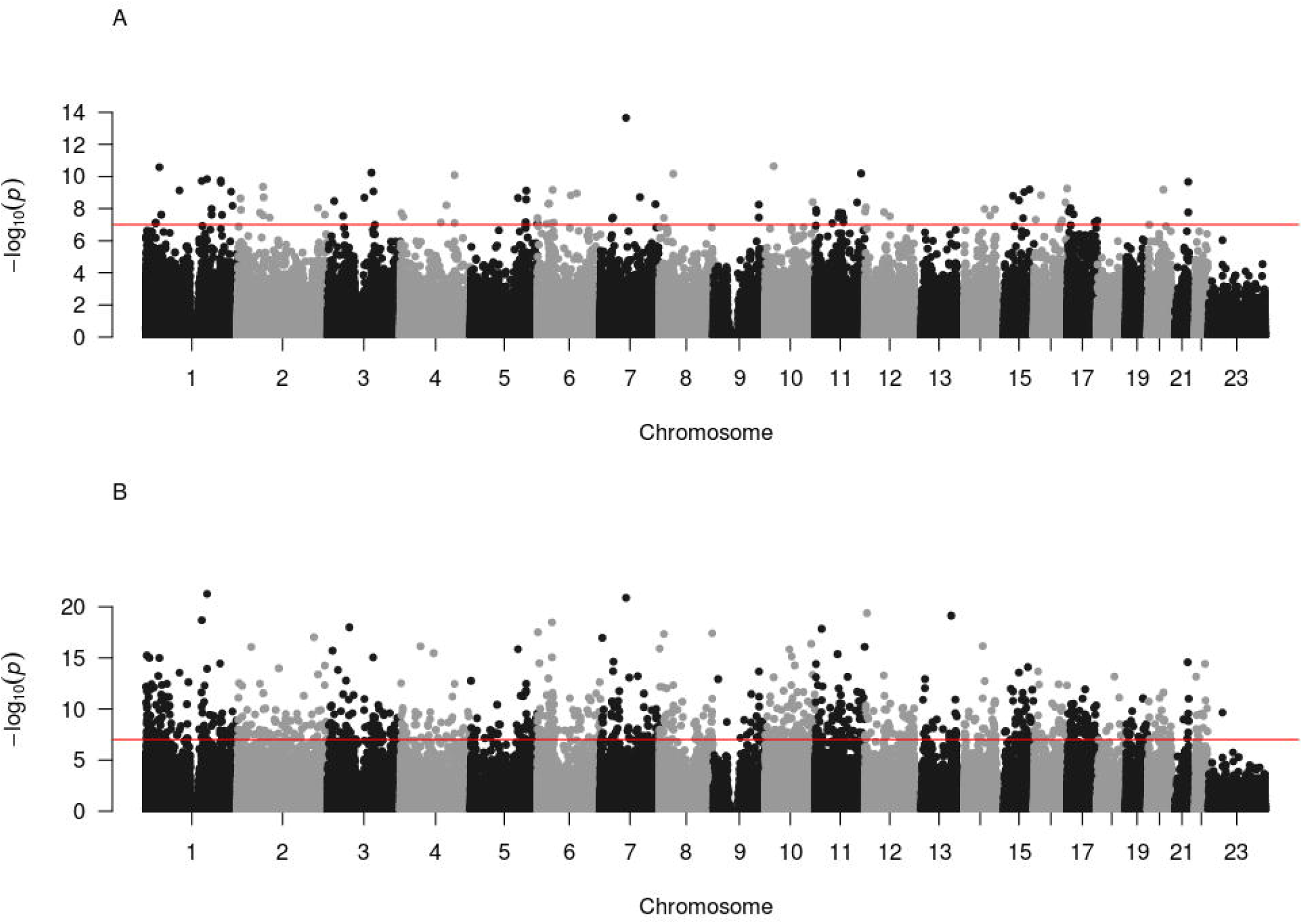
Differential DNA methylation at 93 loci across the genome is associated with psychosis. Manhattan plot depicts the –log10 *P* value from the EWAS meta-analysis (y-axis) against genomic location (x-axis). Panel A) presents results from the analysis comparing psychosis patients and controls, and panel B) presents results from the analysis comparing diagnosed schizophrenia cases and controls.

Because of the importance of considering variation in the composition of the constituent cell types in analyses of complex cellular mixtures ^17,18^, we used established methods to estimate the proportion ^40,41^ and abundance ^38^ of specific cell-types in whole blood. Using a random effects meta-analysis to combine the results across the seven cohorts (**Table 2; Figure 1**), which were adjusted for age, sex and DNAm smoking score, we found that psychosis cases had elevated estimated proportions of granulocytes (mean difference = 0.0431; P = 5.09×10^−4^) and monocytes (mean difference = 0.00320; P = 1.15×10^−4^), and significantly lower proportions of CD4^+^ T-cells (mean difference = -0.0177; P = 0.00144), CD8^+^ T-cells (mean difference = -0.0144; P = 0.00159) and natural killer cells (mean difference = -0.0113; P = 0.00322). Interestingly, the differences in granulocytes, natural killer cells, CD4^+^ T-cells and CD8^+^ T-cells were most apparent in cohorts comprising patients with a diagnosis of schizophrenia (**Figure 1**), with cohorts including FEP patients characterized by weaker or null effects. Limiting the analysis of derived blood cell estimates to a comparison of schizophrenia cases and controls didn’t perceivably change the estimated differences of our random effects model but did reduce the magnitude of heterogeneity compared to including the FEP cases (**eTable 1**). This indicates that changes in blood cell proportions may reflect a consequence of diagnosis, reflecting the fact that people with schizophrenia are likely to have been exposed to a variety of medications, social adversities and somatic ill-health - and for longer periods - than FEP patients. Finally, we used an established algorithm to derive a quantitative DNA methylation “smoking score” for each individual ^42^, building on our previous work demonstrating the utility of this variable for characterizing differences in smoking exposure between schizophrenia patients and controls, and using it as a covariate in an EWAS ^13^. We observed a significantly increased DNA methylation smoking score (**Figure 2**) in psychosis patients compared to controls across all cohorts (mean difference = 3.89; P = 2.88×10^−11^). Although of smaller effect, this difference was also present when comparing FEP and controls in the EU-GEI cohort (mean difference = 2.38; P = 2.68×10^−8^).

**Table 2.** Results of meta-analysis of DNAm estimated cellular composition differences between schizophrenia cases and controls.

| Cell type | Measure type | Number of cohorts | Random effects model |  |  | Fixed effects model |  |  | Heterogeneity P value |
| --- | --- | --- | --- | --- | --- | --- | --- | --- | --- |
|  |  |  | Mean difference | SE | P value | Mean difference | SE | P value |  |
| Monocytes | Proportion | 7 | 0.00312 | 0.00312 | 0.00312 | 0.00312 | 0.00083 | 0.00017 | 0.687 |
| Granulocytes | Proportion | 7 | 0.04360 | 0.04360 | 0.04360 | 0.03952 | 0.00315 | 3.50E-36 | 1.22E-15 |
| Natural Killer cells | Proportion | 7 | -0.01169 | -0.01169 | -0.01169 | -0.00868 | 0.00133 | 6.03E-11 | 1.28E-07 |
| CD4+ T-cells | Proportion | 7 | -0.01728 | -0.01728 | -0.01728 | -0.01509 | 0.00195 | 1.11E-14 | 9.78E-09 |
| CD8+ T-cells | Proportion | 7 | -0.01412 | -0.01412 | -0.01412 | -0.01437 | 0.00149 | 5.18E-22 | 3.23E-10 |
| B-cells | Proportion | 7 | -0.00541 | -0.00541 | -0.00541 | -0.00517 | 0.00102 | 4.10E-07 | 6.95E-06 |
| PlasmaBlast | Abundance | 5 | 0.05362 | 0.05362 | 0.05362 | 0.05366 | 0.00733 | 2.54E-13 | 8.15E-14 |
| CD8pCD28nCD45RAn | Abundance | 5 | 0.06666 | 0.06666 | 0.06666 | 0.11858 | 0.15124 | 0.43303 | 0.091 |
| CD8.naive T-cells | Abundance | 5 | 6.66926 | 6.66926 | 6.66926 | 7.77435 | 1.92790 | 0.00006 | 0.031 |
| CD4.naive T-cells | Abundance | 5 | 9.29613 | 9.29613 | 9.29613 | 9.29613 | 4.76226 | 0.05093 | 0.551 |

### An epigenome-wide association study meta-analysis identifies DNA methylation differences associated with psychosis

To identify differentially methylated positions (DMPs) in blood associated with psychosis, we performed an association analysis within each of the seven schizophrenia and FEP cohorts controlling for age, sex, derived cellular composition variables, derived smoking score, and experimental batch (see **Methods**). We used a Bayesian method to control P-value inflation using the R package *bacon* ^45^ before combining the estimated effect sizes and standard errors across cohorts with a random effects meta-analysis, including all autosomal and X-chromosome DNA methylation sites analyzed in at least two cohorts (n = 839,131 DNA methylation sites) (see **Methods**; **eFigure 4**). Using an experiment-wide significance threshold derived for the Illumina EPIC array ^46^ (P < 9×10^−8^), we identified 95 psychosis-associated DMPs mapping to 93 independent loci and annotated to 68 genes (**Figure 3** and **eTable 2)**. Across these DMPs, the mean difference in DNA methylation between cases and controls was relatively small (0.789%, SD = 0.226%) and there was a striking enrichment of hypermethylated DMPs in psychosis cases (n = 91 DMPs (95.8%) hypermethylated, P = 1.68×10^−22^). A number of the top-ranked DMPs are annotated to genes that have direct relevance to the etiology of psychosis including the GABA transporter *SLC6A12*^56^ (cg00517261, P = 1.53×10^−8^), the GABA receptor *GABBR1*^*^57^*^ (cg00667298, P = 5.07×10^−9^), and the calcium voltage-gated channel subunit gene *CACNA1C* (cg01833890, P = 8.42×10^−9^) that is strongly associated with schizophrenia and bipolar disorder ^58−60^ (**eFigure 5**).

**Figure 4.**
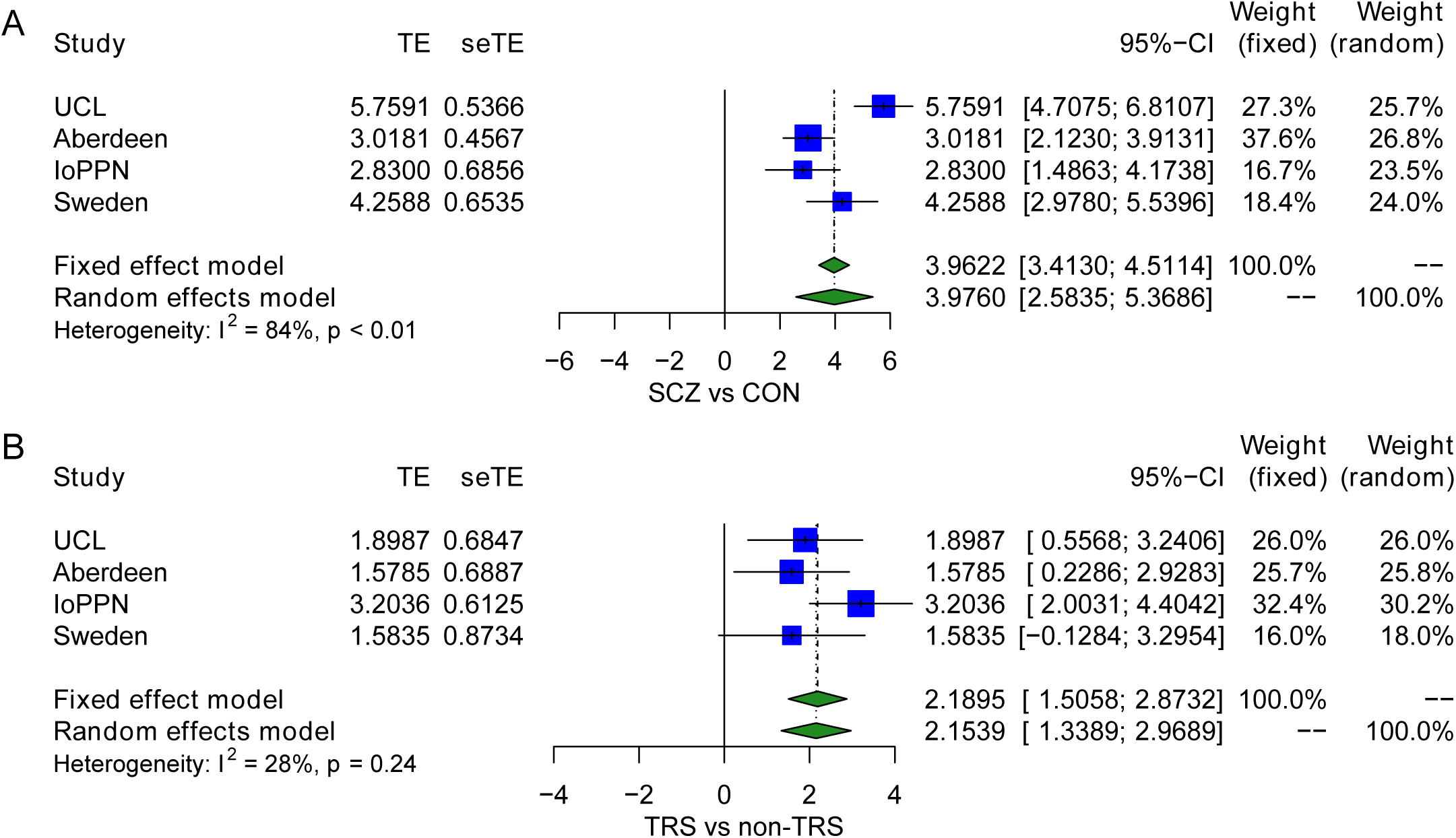
Treatment-resistant schizophrenia patients show an elevated exposure to tobacco smoking relative to non-treatment-resistant schizophrenia and controls. Forest plots from a meta-analysis of differences in smoking score derived from DNA methylation data between **A**) schizophrenia patients and controls and **B**) TRS patients prescribed clozapine and non-TRS prescribed other medications. The smoking score was calculated from DNA methylation data using the method described by Elliott et al^42^. TE – treatment effect i.e. the mean difference between cases and controls, seTE – standard error of the treatment effect.

**Figure 5.**
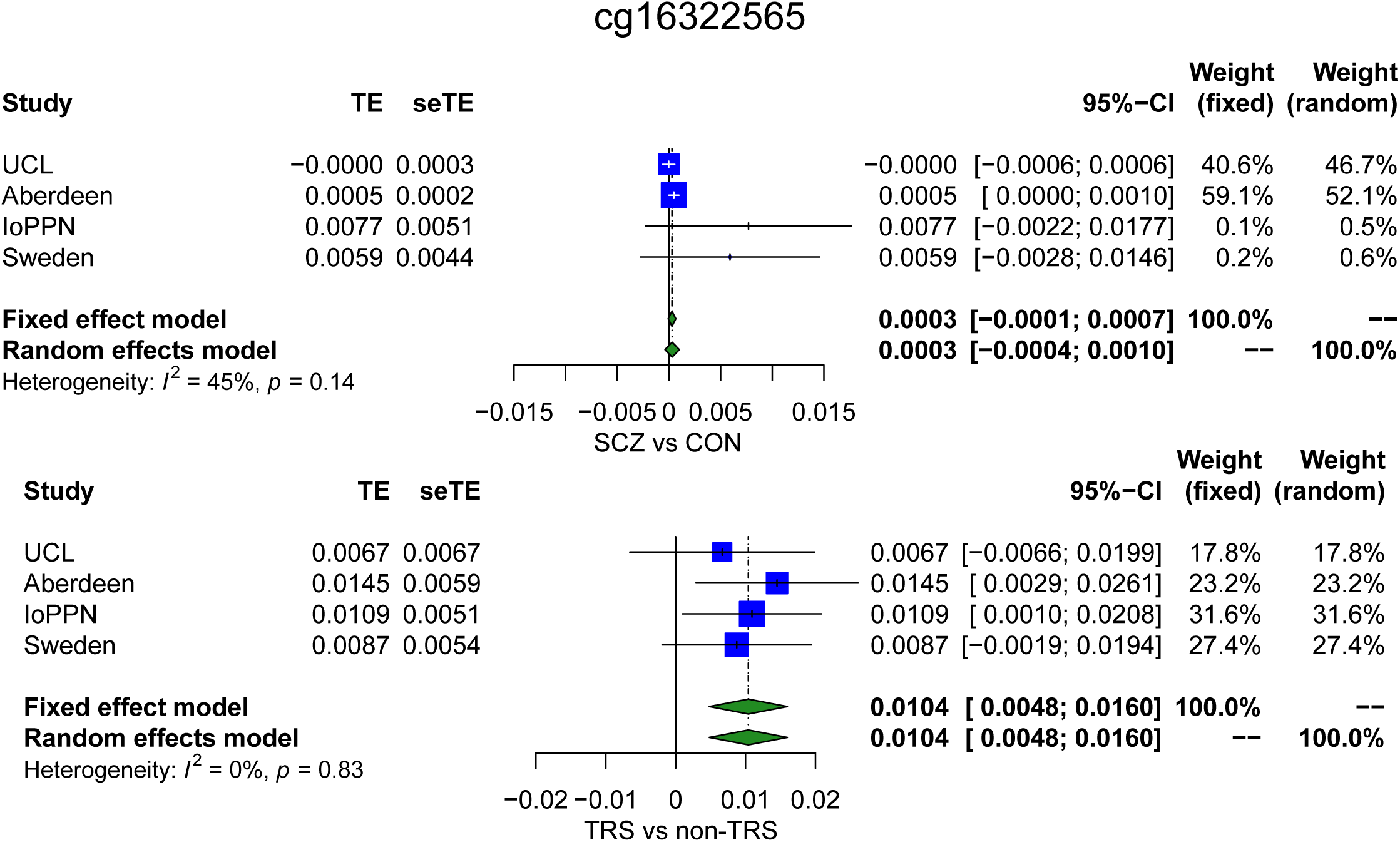
Differences in DNA methylation between schizophrenia cases and controls are driven by subset of cases with treatment resistant schizophrenia. Forest plots from a meta-analysis of differences in DNA methylation at cg16322565 between **A**) schizophrenia patients and controls and **B**) TRS patients prescribed clozapine and non-TRS prescribed other medications. TE – treatment effect i.e. the mean difference between cases and controls, seTE – standard error of the treatment effect.

### A specific focus on clinically-diagnosed schizophrenia cases identifies more widespread DNA methylation differences

We next repeated the EWAS focussing specifically on the subset of psychosis cases with diagnosed schizophrenia (schizophrenia cases = 1,681, controls = 1,583; **eFigure 4**). Compared to our EWAS of psychosis we identified more widespread differences in DNA methylation, with 1,048 schizophrenia associated DMPs (P < 9×10^−8^) representing 1,013 loci and annotated to 692 genes (**eTable 3**).

Although the list of schizophrenia-associated DMPs included 61 (64.21%) of the psychosis associated DMPs, the total number of significant differences was much larger, potentially reflecting the less heterogeneous clinical characteristics of the cases. Schizophrenia-associated DMPs had a mean difference of 0.789% (SD = 0.204%), and like the psychosis-associated differences, were significantly enriched for sites that were hypermethylated in cases compared to controls (n = 897, 87.4%, P = 1.27×10^−129^)). A number of the top-ranked DMPs are annotated to genes that have direct relevance to the etiology of schizophrenia and gene ontology (GO) analysis highlighted multiple pathways previously implicated in schizophrenia including several related to the extracellular matrix^61^ and cell-cell adhesion^62^ (**eTable 4**).

### Schizophrenia-associated DMPs colocalize to regions nominated by genetic association studies

As the etiology of schizophrenia has a large genetic component, we next sought to explore the extent to which DNA methylation at schizophrenia-associated DMPs is influenced by genetic variation. Using results from a quantitative genetic analysis of DNA methylation in monozygotic and dizygotic twins ^63^, we found that DNA methylation at schizophrenia-associated DMPs is more strongly influenced by additive genetic factors compared to non-associated sites matched for comparable means and standard deviations (**eFigure 6**) (mean additive genetic component across DMPs = 23.0%; SD = 16.8%; P = 1.61×10^−87^; **eFigure 7**). Using a database of blood DNA methylation quantitative trait loci (mQTL) previously generated by our group ^12^ we identified common genetic variants associated with 256 (24.4%) of the schizophrenia-associated DMPs. Across these 256 schizophrenia-associated DMPs there were a total of 455 independent genetic associations with 448 genetic variants, indicating that some of these DMPs are under polygenic control with multiple genetic variants associated. Of note, 31 of these genetic variants are located within 12 schizophrenia-associated GWAS regions (**eTable 5**) with 19 genetic variants associated with schizophrenia DMPs located in the MHC region on chromosome 6. To further support an overlap between GWAS and EWAS signals for schizophrenia, we compared the list of genes identified in this study with those from the largest GWAS meta-analysis of schizophrenia ^6^ identifying 21 schizophrenia-associated DMPs located in 11 different GWAS regions. To more formally test for an enrichment of differentially methylation across schizophrenia-associated GWAS regions, we calculated a combined EWAS P-value for each of the GWAS associated regions using all DNA methylation sites within each region identifying 21 significant regions (P < 3.16×10^−4^, corrected for testing 158 regions; **eTable 6**). Three of these regions also contained a significant schizophrenia-associated DMP and a genetic variant associated with that schizophrenia-associated DMP. These include a region located within the MHC, another located on chromosome 17 containing *DLG2, TOM1L2* and overlapping the Smith-Magenis syndrome deletion, and another on chromosome 16 containing *CENPT*, and *PRMT7*.

### Treatment-resistant schizophrenia cases differ from treatment-responsive schizophrenia patients for blood cell proportion estimates and smoking score derived from DNA methylation data

Up to 25% of schizophrenia patients are resistant to the most commonly prescribed antipsychotic medications, and clozapine is a second-generation antipsychotic often prescribed to patients with such treatment-resistant schizophrenia (TRS) who may represent a more severe subgroup ^64^. Using data from four cohorts for which medication records were available (UCL, Aberdeen, IoPPN, and Sweden), we performed a within-schizophrenia analysis comparing schizophrenia patients prescribed clozapine (described as TRS cases) and those prescribed standard antipsychotic medications (total n = 399 TRS and 636 non-TRS). Across each of the four cohorts the proportion of males prescribed clozapine was slightly higher than the proportion of males on other medications (P = 0.0211; **eTable 7**) consistent with findings from epidemiological studies that report increased rates of clozapine prescription in males^65^, although there was statistically significant heterogeneity in the sex distribution between groups across cohorts (P = 7.96×10^−3^). There was no overall significant difference in age between TRS and non-TRS cases (P = 0.533; **eFigure 8**), although there was significant heterogeneity between the cohorts (P = 7.40×10^−32^). There was no evidence of accelerated epigenetic aging between TRS and non-TRS patients (**eFigure 9** and **eFigure 10**). Interestingly, cellular composition variables derived from the DNA methylation data suggests that TRS cases are characterized by a significantly higher proportion of granulocytes (meta-analysis mean difference = 0.00283; P = 8.10×10^−6^) and lower proportions of CD8^+^ T-cells (mean difference = -0.0115; P = 4.37×10^−5^ (**eTable 8; eFigure 11**) compared to non-TRS cases. Given the finding of higher derived granulocyte and lower CD8^+^ T-cell levels in the combined psychosis patient group compared to controls described above, a finding driven primarily by patients with schizophrenia, we performed a multiple regression analysis of granulocyte proportion to partition the effects associated with schizophrenia status from effects associated with TRS status. After including a covariate for TRS, schizophrenia status was not significantly associated with granulocyte proportion using a random effects model (P = 0.210) but there was significant heterogeneity of effects across the four cohorts (P = 4.93×10^−7^). Within the group of patients with schizophrenia, however, there were notable differences between TRS and non-TRS groups (mean difference = 0.0275; P = 3.22×10^−6^; **eFigure 12)**. In contrast a multiple regression analysis found that both schizophrenia status (mean difference = -0.0113; P = 0.00818) and TRS status (mean difference = -0.0116; P = 2.82×10^−5^) had independent additive effects on CD8^+^ T-cell proportion (**eFigure 13**). Finally, TRS was also associated with significantly higher DNA methylation-derived smoking scores than non-TRS in all four cohorts (mean difference = 2.16; P = 7.79×10^−5^; **eFigure 14**). Testing both schizophrenia diagnosis status and TRS status simultaneously, we found that both remained significant; schizophrenia diagnosis was associated with a significant increase in smoking score (P = 2.19×10^−8^) with TRS status associated with an additional increase within cases (P = 2.22×10^−7^) (**Figure 4**).

### There are widespread DMPs between treatment-resistant schizophrenia patients and treatment-responsive patients

We next performed an EWAS within schizophrenia patients comparing TRS cases to non-TRS cases, including each autosomal and X-chromosome DNA methylation site analyzed in at least two cohorts (n = 431,659 DNA methylation sites). We identified seven DMPs associated with clozapine exposure (P < 9×10^−8^; **eTable 9**) with a mean difference of 1.47% (SD = 0.242%) and all sites being characterized by elevated DNA methylation in TRS cases (P = 0.0156). We were interested in whether the DNA methylation differences associated with TRS overlapped with those identified between all schizophrenia cases and non-psychiatric controls. Although there was no direct overlap between the clozapine associated DMPs and the schizophrenia associated DMPs identified for each analysis, the direction of effects across the 1,048 schizophrenia-associated DMPs were strikingly consistent (n = 738 (70.4%) DMPs with consistent direction; P = 7.57×10^−41^; **eFigure 15**). Given these observations, we formally tested whether the schizophrenia-associated differences are driven by the subset of TRS cases on clozapine by fitting a model that simultaneously estimates the effect of schizophrenia status and TRS status across all 1,048 sites (**eTable 10**). While the vast majority of schizophrenia associated DMPs remained at least nominally significant (n = 1,003 95.7%, P < 0.05) between schizophrenia patients and controls, amongst those that didn’t 25 (2.39%) had a significant effect associated with TRS status. For example, differential DNA methylation at the schizophrenia-associated DMP cg16322565, located in the NR1L2 gene on chromosome 3 (schizophrenia EWAS meta-analysis: mean DNA methylation difference = 0.907%, P = 3.52×10^−9^), is driven primarily by cases with TRS (**Figure 5**; multivariate analysis mean DNA methylation difference between schizophrenia cases and controls = 0.323%, P = 0.123, mean DNA methylation difference between TRS cases and non-TRS controls = 1.01%, P = 8.71×10^−5^). 152 (14.5%) of the schizophrenia associated DMPs were associated with a significant effect between schizophrenia cases and controls and a significant affect within schizophrenia patients between TRS and non-TRS patients, with the majority (128 (84.2%)) characterized by the same direction of effect in both groups and indicative of an additive effect of both schizophrenia diagnosis and TRS status (e.g. **eFigure 16**). Of particular interest are 24 DMPs which are significantly associated with both schizophrenia and TRS but with an opposite direction of effect, highlighting how that at some DNA methylation sites, TRS counteracts changes induced by schizophrenia (e.g. **eFigure 17)**. Taken together, 177 (16.9%) of the schizophrenia-associated DMPs identified in our EWAS meta-analysis are influenced by TRS and reflect either differences induced by exposure to a specific antipsychotic therapy or other differences (e.g. treatment resistance) in individuals who are prescribed clozapine.

## Discussion

We report a comprehensive study of methylomic variation associated with psychosis and schizophrenia, profiling DNA methylation across the genome in peripheral blood samples from 2,379 cases and 2,104 controls. We show how DNA methylation data can be leveraged to derive measures of blood cell counts and smoking that are associated with psychosis. Using a stringent pipeline to meta-analyze EWAS results across datasets, we identify DMPs associated with both psychosis and a more refined diagnosis of schizophrenia. Of note, we show evidence for the co-localization of genetic associations for schizophrenia and differential DNA methylation. Finally, we present evidence for differential methylation associated with treatment-resistant schizophrenia, potentially reflecting exposure to clozapine.

We identify psychosis-associated differences in cellular composition estimates derived from DNA methylation data, with cases having increased proportions of monocytes and granulocytes and decreased proportions of natural killer cells, CD4^+^ T-cells and CD8^+^ T-cells compared to non-psychiatric controls. This analysis extends previous work based on a subset of these data, which reported a decrease in the proportion of natural killer cells and increase in the proportion of granulocytes in schizophrenia patients^29^, with the large number of samples enabling us to identify additional associations with other cell types. We also confirm findings from an independent study of schizophrenia which reported significantly increased proportions of granulocytes and monocytes, and decreased proportions of CD8^+^ T-cells using estimates derived from DNA methylation data ^66^. Of note, because we can only derive proportion of cell types from whole blood DNA methylation data, and not actual counts, an increase in one or more cell types must be balanced by a decrease in one or more other cell types and an apparrent change in the proportion of one specific cell type does not mean that the actual abundance of that cell type is altered. Despite this, the results from DNA methylation-derived cell proportions are consistent with previous studies based on empirical cell abundance measures which have reported increased monocyte counts^67,68^, increased neutrophil counts^69,70^, increased monocyte to lymphocyte ratio^71,72^ and increased neutrophil to lymphocyte ratio ^71,73^ in both schizophrenia and FEP patients compared to controls. Studies have also shown that higher neutrophil counts in schizophrenia patients correlate with a greater burden of positive symptoms^69^ suggesting that variations in the number of neutrophils is a potential marker of disease severity^72^. Our sub-analysis of treatment-resistant schizophrenia, which is associated with a higher number of positive symptoms ^65^, found that the increase in granulocytes was primary driven by those with the more severe phenotype, supporting this hypothesis. Importantly, the differences we observe may actually reflect the effects of various antipsychotic medications that have been previously shown to influence cell proportions in blood^72^.

We also identified a significant increase in a DNA methylation-derived smoking score in patients with schizophrenia, replicating our previous finding ^13^. The smoking score captures multiple aspects of tobacco smoking behaviour including both current smoking status and the quantity of cigarettes, smoked; our results therefore reflect existing epidemiological evidence demonstrating that schizophrenia patients not only smoke more, but also smoke more heavily (55-57). We also report an increased smoking score in patients with FEP, albeit with of smaller magnitude than seen in schizophrenia, consistent with a meta-analysis reporting increasing levels of smoking in FEP (58). In the subset of treatment-resistant patients, we found that there was an additional increase in smoking score relative to schizophrenia cases prescribed alternative medications, supporting evidence for higher rates of smoking in TRS groups relative to treatment-responsive schizophrenia patients^74^. These results not only highlight important biological and environmental differences associated with psychosis and schizophrenia, but also highlight the importance of controlling for these differences as potential confounders in studies of disease-associated DNA methylation differences.

Our epigenome-wide association study, building on our previous analysis on a subset of the sample cohorts profiled here ^13^ identified 95 DMPs associated with psychosis that are robust to differences in measured smoking exposure and heterogeneity in blood cellular composition derived from DNA methylation data. Of note, we identified a dramatic increase in sites characterized by an increase in DNA methylation in patients. A key strength of our study is the inclusion of the full spectrum of schizophrenia diagnoses, from FEP through to treatment-resistant cases prescribed clozapine. While this may introduce heterogeneity into our primary analyses, we used a random effects meta-analysis to identify consistent effects across all cohorts and diagnostic subtypes. We also performed an additional analysis focused specifically on cases with diagnosed schizophrenia excluding those with FEP, which identified many more DMPs. Our results suggest that this analysis of a more specific phenotype in a smaller number of samples is potentially more powerful and that schizophrenia cases have a more discrete molecular phenotype that might reflect both etiological factors but also factors associated with a diagnosis of schizophrenia (e.g. medications, stress, etc). The mean difference in DNA methylation between cases and controls for both psychosis and schizophrenia was small, consistent with other blood-based EWAS of schizophrenia ^66^ and complex traits ^75-77^ in general. While individually they may be too small to have a strong predictive power as a biomarker, together they may have utility as a molecular classifier^78^.

We also report the first systematic analysis of individuals with TRS, identifying seven DMPs at which differential DNA methylation was significantly different in the subset of schizophrenia cases prescribed clozapine. These data are highly informative for the interpretation of our schizophrenia-associated differences, because a number of these DMPs are driven by the subset of patients on clozapine. Furthermore, a number of sites show opposite effects in our analyses of TRS vs our analysis of schizophrenia, suggesting they might represent important differences between diagnostic groups. Because the prescription of clozapine is generally only undertaken in patients with treatment-resistant schizophrenia, we are unable to separate the effects of clozapine exposure from differences associated with a more severe sub-type of schizophrenia.

Our results should be considered in light of a number of important limitations. First, our analyses were constrained by the technical limitations of the Illumina 450K and EPIC arrays which only assays ∼ 3% of CpG sites in the genome. Second, this is a cross-sectional study and was not possible to distinguish cause from effect. It is possible, and indeed likely, for example, that the differences associated with both schizophrenia and TRS reflect the effects of medication exposure or other consequences of having schizophrenia, e.g. living more stressful lives, poorer diet and health. The importance of such confounding variables is demonstrated by our findings of differential smoking score and blood cell proportions derived directly from the DNA methylation data, although these examples also highlight the potential utility of such effects for molecular epidemiology. Third, this work is based on DNA methylation profiled in a peripheral blood and therefore can provide only limited information about variation in the brain^79^. This is a salient point for understanding the role DNA methylation plays in the disease process, but biomarkers, by definition, need to be measured in an accessible tissue and don’t necessarily need to reflect the underlying pathogenic process. Furthermore, because most classifiers used to quantify variables such as smoking exposure and age have been trained in blood, this represents the optimal tissue in which to derive these measures. Of course, blood may also be an appropriate choice for investigating medication effects, particularly given the known effects on white blood cell counts associated with taking clozapine^80^. Fourth, while we have explored the potential effects of clozapine on DNA methylation by assessing a sub-group of individuals with TRS, this is just one of a range of antipsychotics schizophrenia and psychosis patients are prescribed. The fact that the TRS group show more extreme differences for many of the schizophrenia-associated DMPs suggests that the polypharmaceutical treatment regimes often prescribed to schizophrenia patients may produce specific DNA methylation signatures in patients, akin to the effect seen for smoking.

In conclusion, we report the largest study of blood based DNA-methylation in schizophrenia and psychosis, and the first within case analysis of treatment-resistant schizophrenia. Our results highlight differences in blood cellular composition and smoking behaviour between not just cases and controls, but also between treatment-resistant schizophrenia patients prescribed clozapine and those prescribed alternative medications. We report widespread differences in DNA methylation in psychosis and schizophrenia, a subset of which are driven by the more severe treatment-resistant subset.

## Authors’ contributions

JM obtained funding and supervised the project. EH lead and performed the analysis with support from ED and GM. ED and JB undertook laboratory work. EH and JM drafted the manuscript. GB, DC, RM and LSS were co-applicants on funding application. AC, CJC, DD MDF, TGD, GD, FG, MG, AG, CG, HEH, CMH, VJ, RSK, JK, GK, KK, JMac, AM, CM, DWM, KCM, CM, IN, MCO, DQ, ALR, BPFR, DSC, eTable, TT, JVO, JLW, PFS, GB, RM were involved in recruiting the samples, providing extracted DNA samples for processing, and collecting the phenotype information. All authors read and approved the final manuscript.

## Acknowledgments

This work was primarily supported by grants from the UK Medical Research Council (MRC; MR/K013807/1 and MR/R005176/1) to J.M. High-performance computing was supported by MRC Clinical Research Infrastructure Funding (MR/M008924/1). The Finnish Twin study was supported by the Academy of Finland Centre of Excellence in Complex Disease Genetics (grant numbers: 213506, 129680), and J.K. by the Academy of Finland grants 265240, 263278 and 312073. Financial support for the Sweden twin study was provided by the Karolinska Institutet (ALF 20090183 and ALF 20100305 to Hultman) and NIH (R01 MH52857). Collection of the Sweden case control samples was supported by the Sweden Research Council (Vetenskapsrådet, award D0886501 to PFS) and the NIMH (R01MH077139). Collection of the Irish case control samples was funded by the Wellcome Trust Case Control Consortium 2 project (085475/B/08/Z and 085475/Z/08/Z), the Wellcome Trust (072894/Z/03/Z, 090532/Z/09/Z and 075491/Z/04/B), and Science Foundation Ireland (08/IN.1/B1916). The European Network of National Schizophrenia Networks Studying Gene-Environment Interactions (EU-GEI) Project is funded by grant agreement HEALTH-F2-2010-241909 (Project EU-GEI) from the European Community’s Seventh Framework Programme. The IMPaCT programme at King’s College London and the South London and Maudsley NHS Foundation Trust is funded by the National Institute for Health Research (RP-PG-0606-1049). The CREeTable AR project received funding from the European Union’s Seventh Framework Programme for research, technological development and demonstration under grant agreement 279227 (CREeTable AR Consortium). EH, ED, LS and JM were supported by MRC grant K013807 to JM. Cardiff University researchers were supported by Medical Research Council (MRC) Centre (G0800509) and Programme Grant (G0801418). Bart PF Rutten is supported by a VIDI grant (number 91718336) from the Netherlands Organisation for Scientific Research. FG is in part supported by the National Institute for Health Research’s (NIHR) Biomedical Research Centre at South London and Maudsley NHS Foundation Trust and King’s College London, the Stanley Medical Research Institute, the Maudsley Charity and the National Institute for Health Research (NIHR) Applied Research Collaboration South London (NIHR ARC South London) at King’s College Hospital NHS Foundation Trust. MDF and DQ are funded by an MRC fellowship to MDF (MR/M008436/1). We gratefully acknowledge capital equipment funding from the Maudsley Charity (Grant Ref. 980) and Guy’s and St Thomas’s Charity (Grant Ref. eTable R130505). This study presents independent research supported by the National Institute for Health Research NIHR BioResource Centre Maudsley at South London and Maudsley NHS Foundation Trust and King’s College London. The views expressed are those of the author(s) and not necessarily those of the NHS, NIHR, Department of Health and Social Care or King’s College London.

