## Supplementary Figures for "Large-scale analysis of DNA methylation identifies cellular alterations in blood from psychosis patients and molecular biomarkers of treatment-resistant schizophrenia"

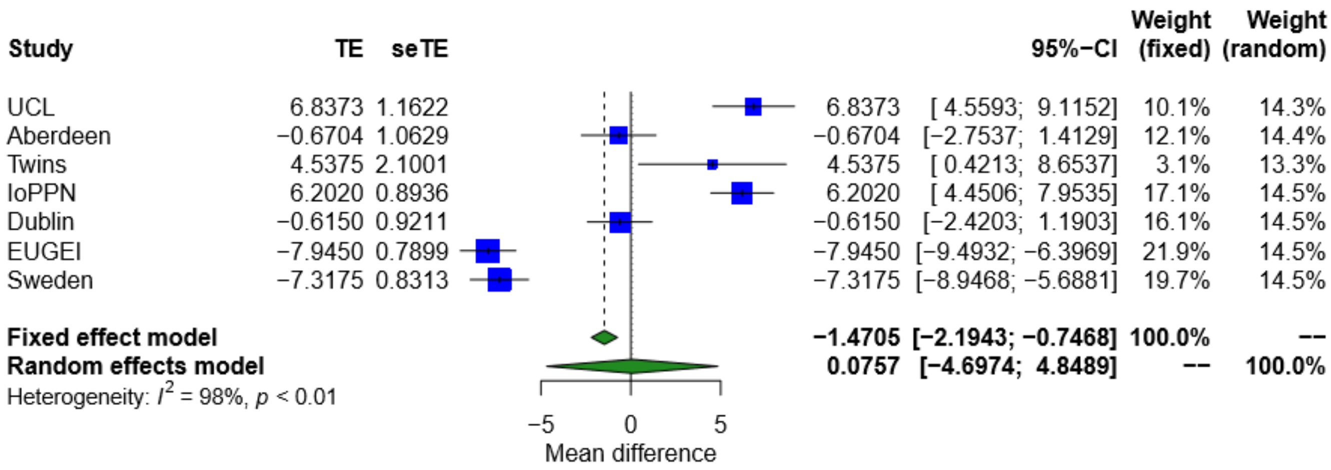


**SF1: Forest plot showing the difference in mean age between psychosis cases and controls across each cohort.** TE – treatment effect i.e. the mean difference between cases and controls, seTE – standard error of the treatment effect.

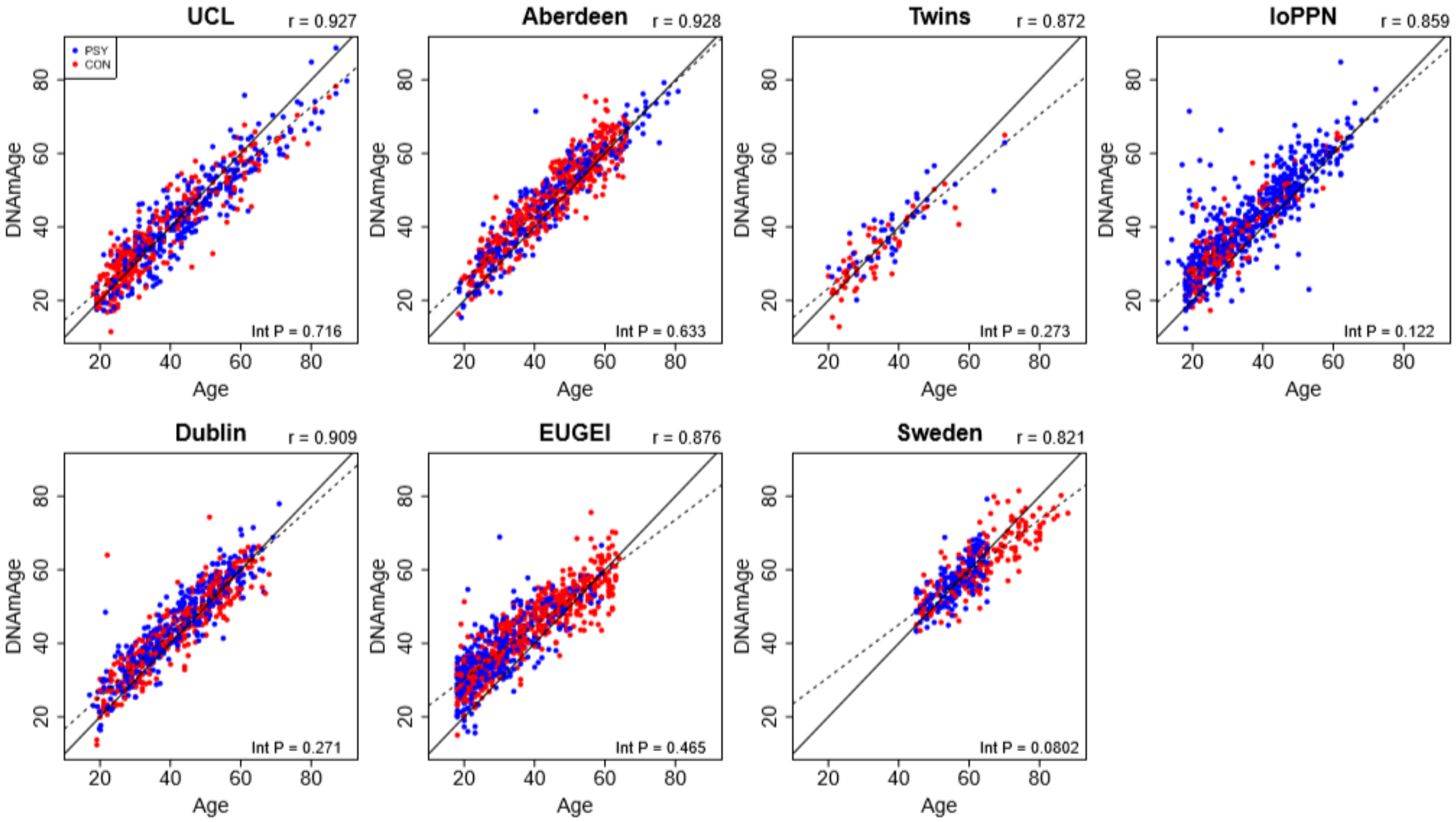


**SF2: Scatterplots of DNAmAge derived from the DNA methylation data against actual chronological age for each of the cohorts.** DNAmAge was calculated using the algorithm described by Horvath et al[1]. Each point represents an individual and is coloured by psychosis status (blue = psychosis, red = control). The solid diagonal line depicts x=y, i.e. where the estimated and actual values are the same. The dashed diagonal line depicts the line of best fit. Presented at the top of the graph is the Pearson's correlation coefficient (r) between the estimated and actual age across all samples in that cohort. Also shown in the bottom right hand corner of each panel is an interaction P value from a test for different correlations between DNAmAge and actual age between psychosis cases and controls.

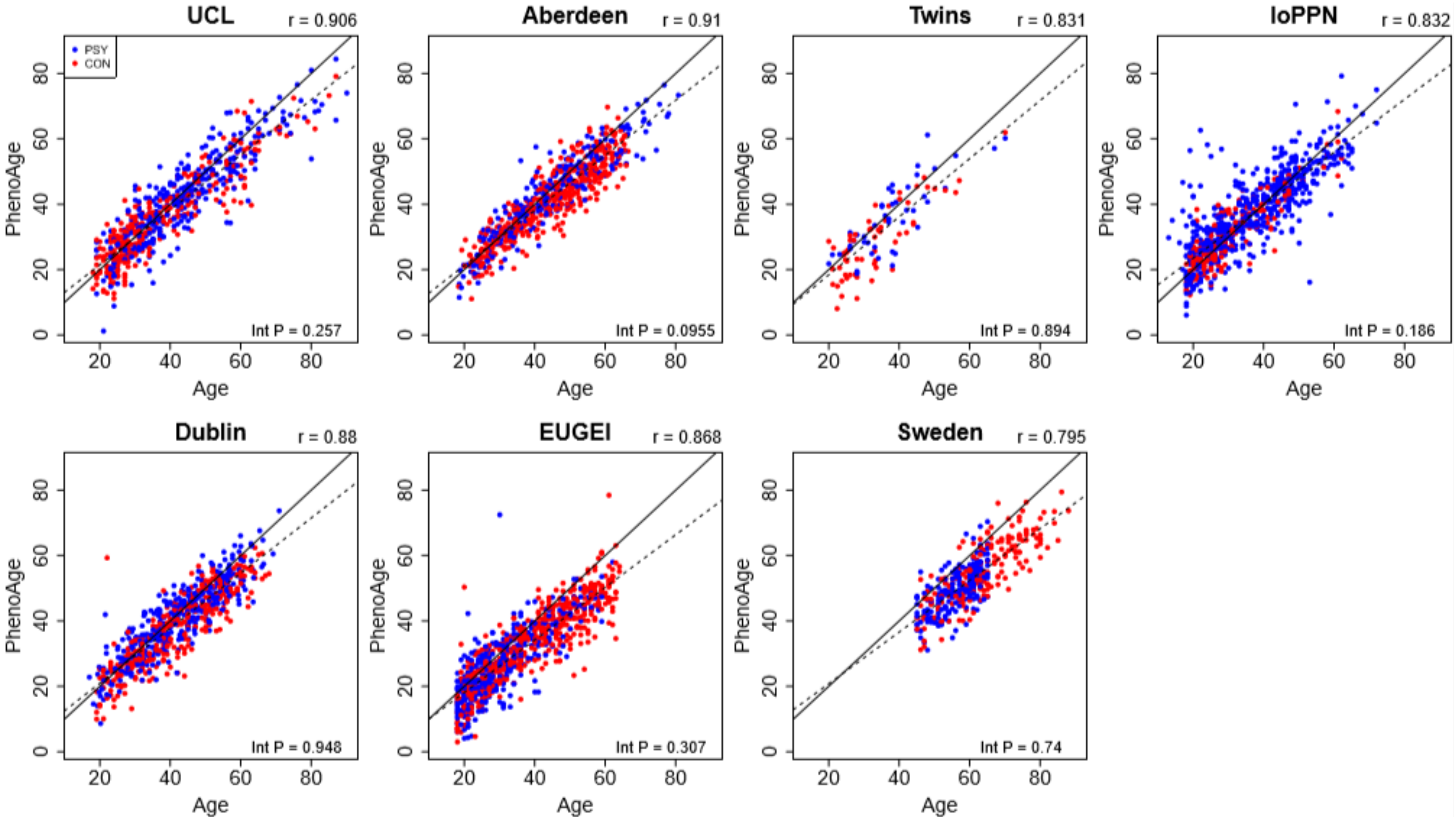


**SF3: Scatterplots of PhenoAge derived from DNA methylationdata against actual chronological age for each of the cohorts.** PhenoAge was calculated using the algorithm described by Levine et al. [1]. Each point represents an individual and is coloured by psychosis status (blue = psychosis, red = control). The solid diagonal line depicts x=y, i.e. where the estimated and actual values are the same. The dashed diagonal line depicts the line of best fit. Presented at the top of the graph is the Pearson's correlation coefficient (r) between the estimated and actual age across all samples in that cohort. Also shown in the bottom right hand corner of each panel is an interaction P value from a test for different correlations between PhenoAge and actual age between psychosis cases and controls.

A

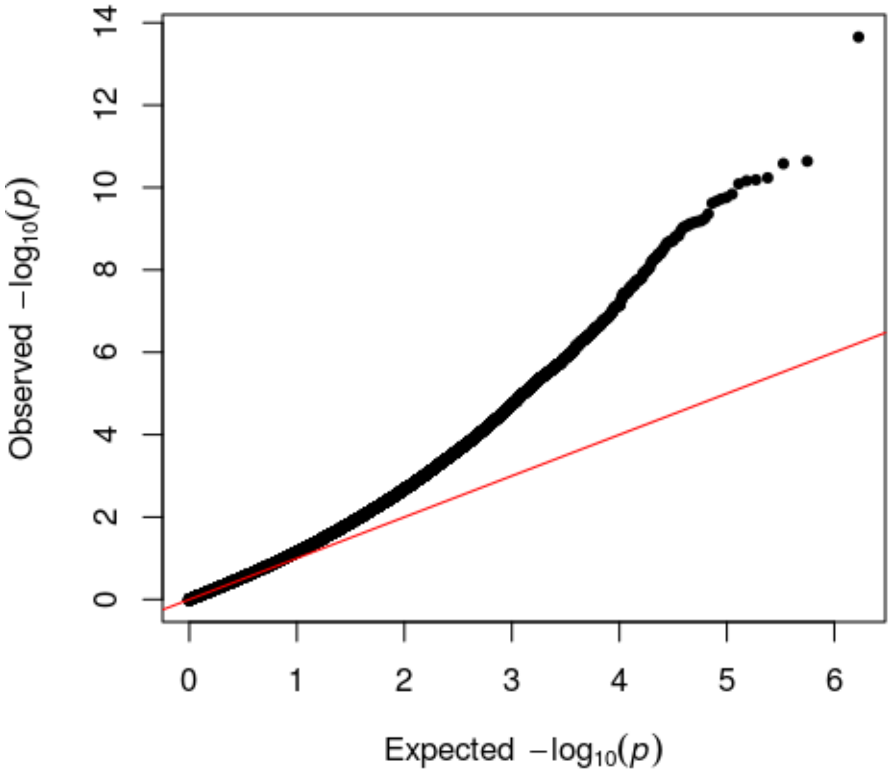


B

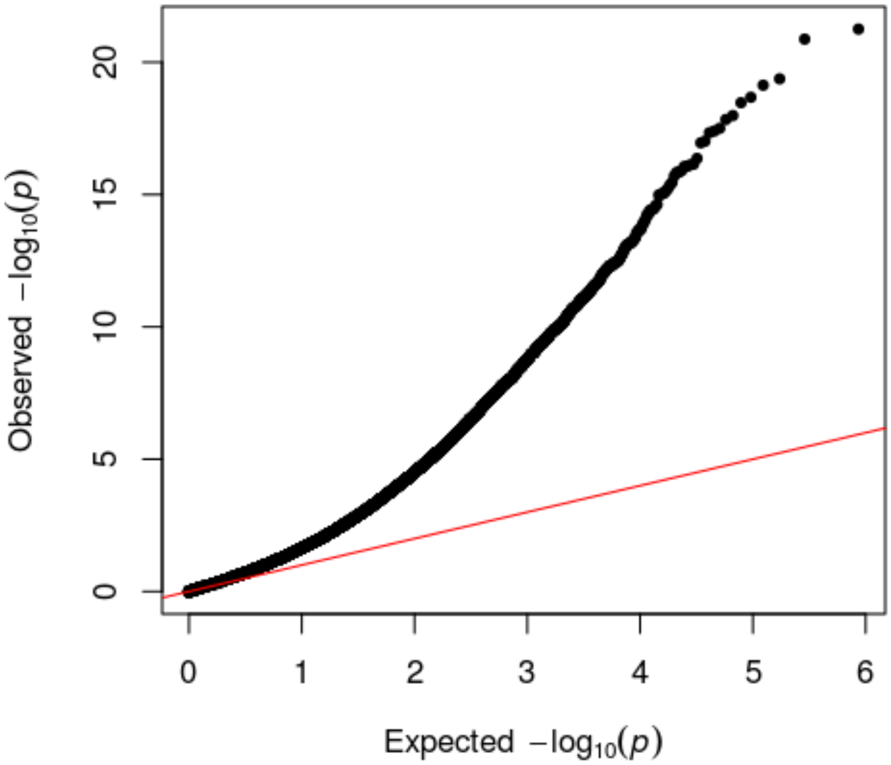


**SF4: QQ plot of EWAS in whole blood**. QQ plots of p values from random effects meta-analysis of A) psychosis cases vs controls and B) schizophrenia cases vs controls.

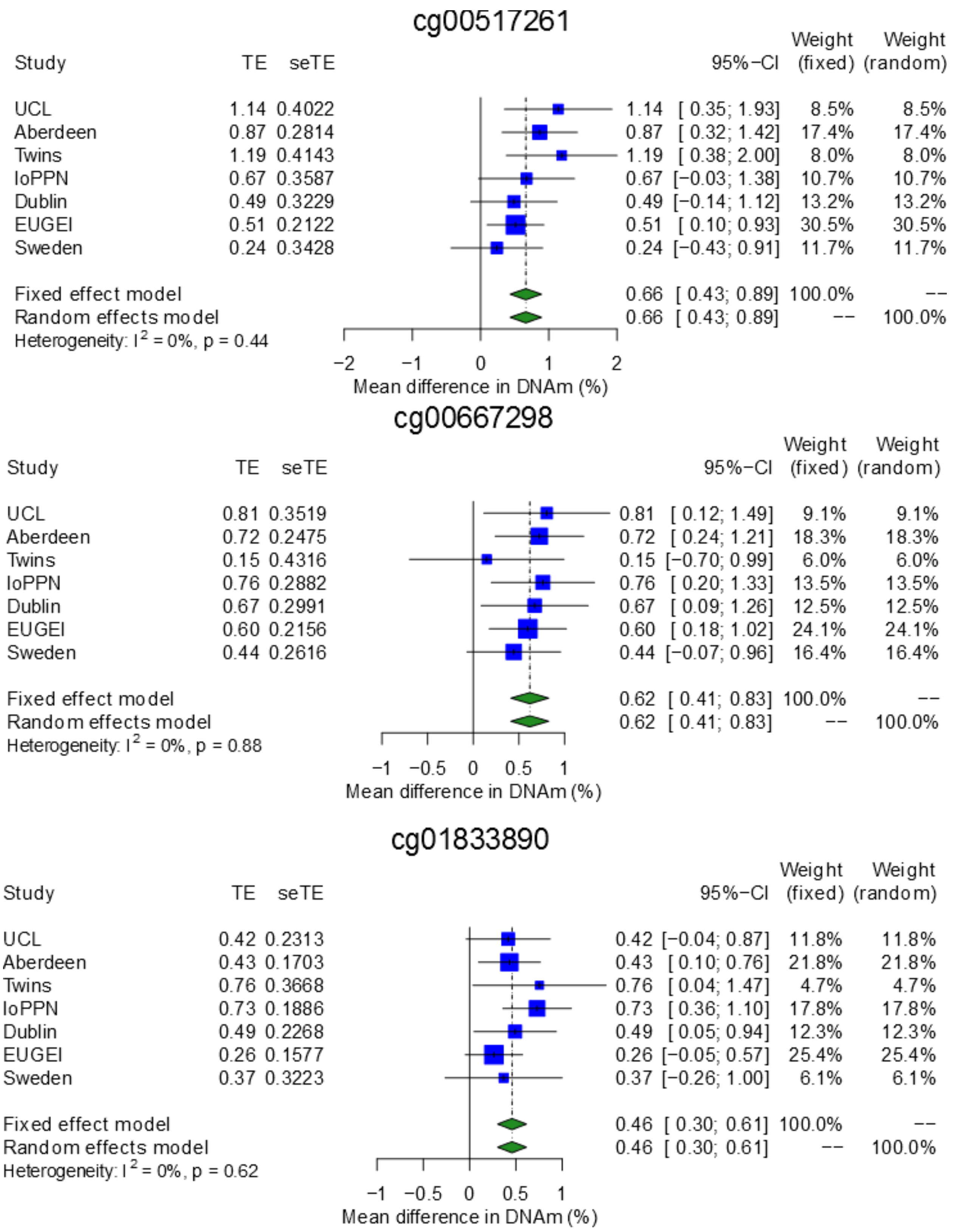


**SF5: Forest plots of notable psychosis associated differentially methylated positions.** TE – treatment effect i.e. the mean difference between cases and controls, seTE – standard error of the treatment effect.

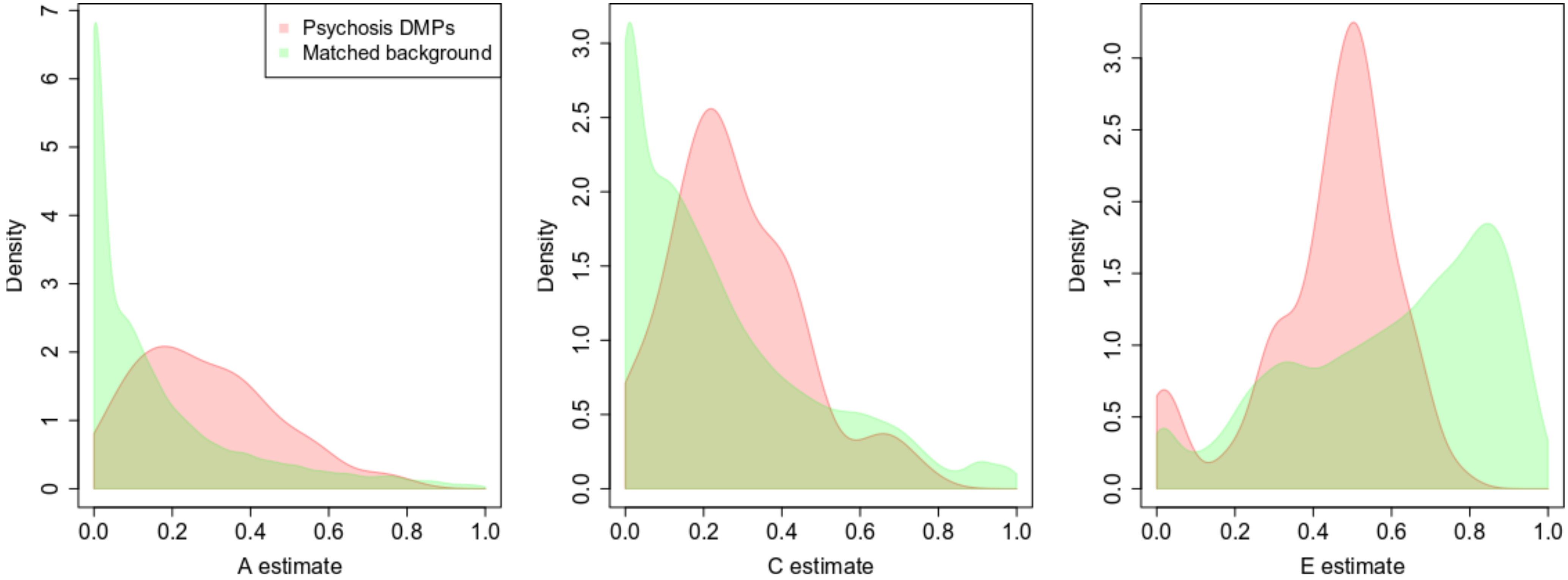


**SF6. DNA methylation at sites associated with psychosis is more strongly influenced by genetic factors and common environmental influences than equivalent matched sites across the genome.** A series of density plots for estimates of additive genetic effects (A, left), common environmental effects (C, middle), and non-shared environmental effects (E, right) derived using data from Hannon et al [2].

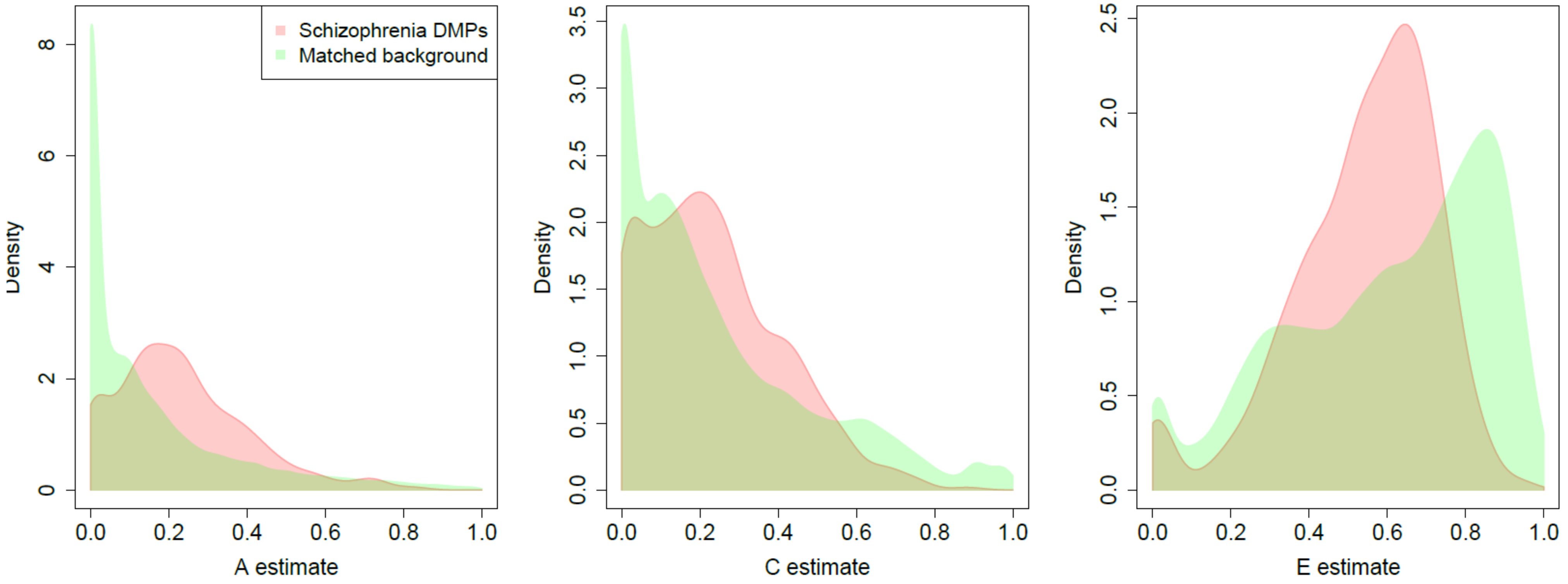


**SF7. DNA methylation at sites associated with schizophrenia is more strongly influenced by genetic factors and common environmental influences than equivalent matched sites across the genome.** A series of density plots for estimates of additive genetic effects (A, left), common environmental effects (C, middle), and non-shared environmental effects (E, right) derived using data from Hannon et al [2].

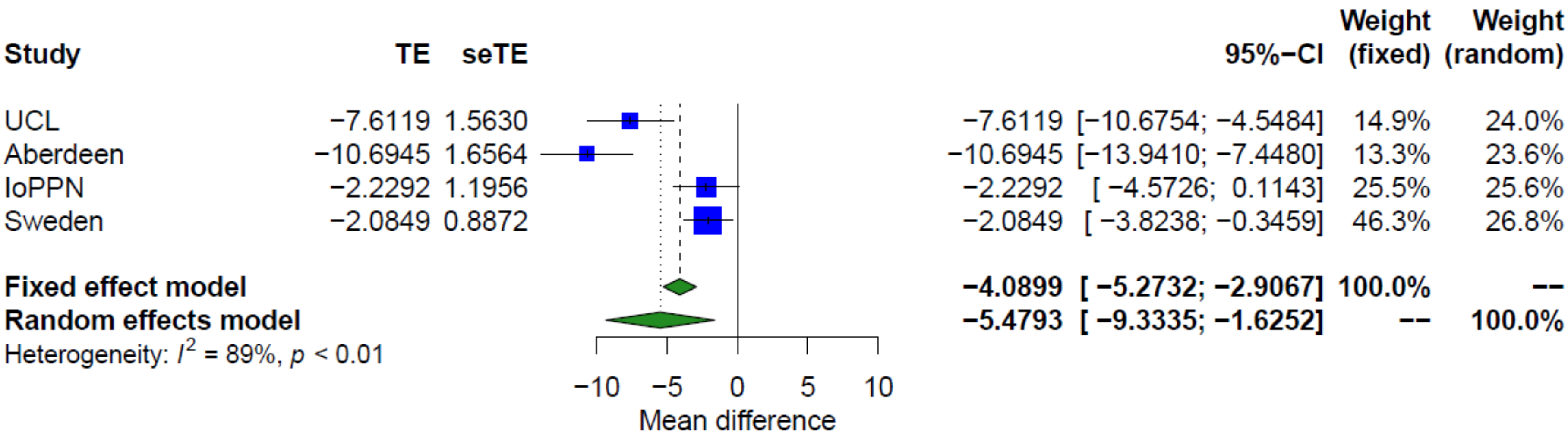


**SF8. Forest plot showing the difference in age between treatment-resistant schizophrenia cases prescribed clozapine and schizophrenia cases on alternative medications across each cohort.** TE – treatment effect i.e. the mean difference between cases and controls, seTE – standard error of the treatment effect.

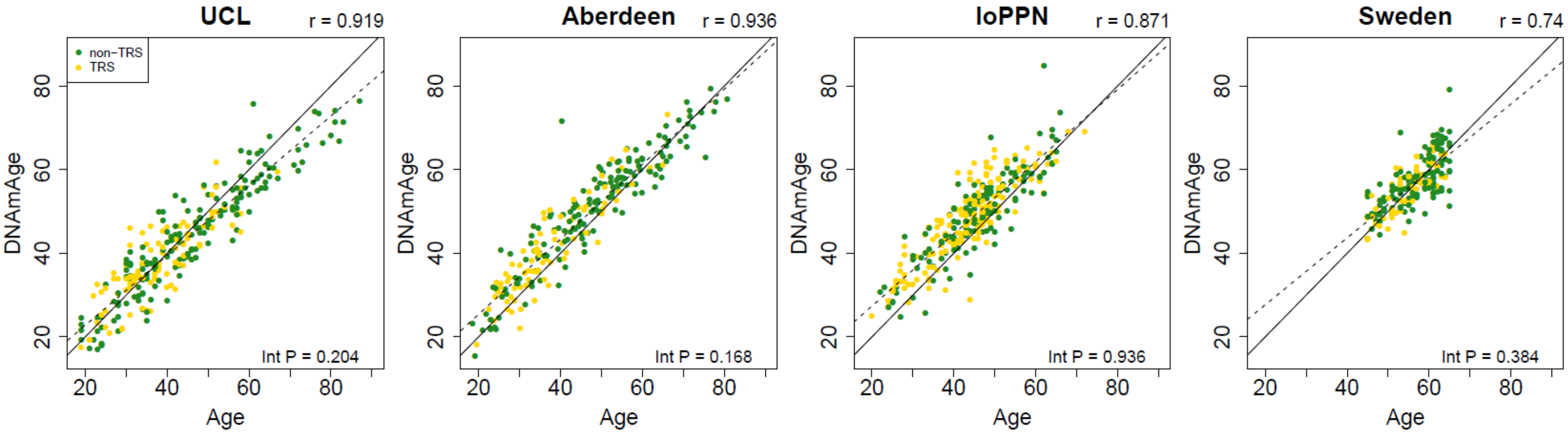


**SF9:** Scatterplots of DNAmAge derived from the DNA methylation data against actual chronological age for each of the cohorts. DNAmAge was calculated using the algorithm described by Horvath et al[3]. Each point represents an individual and is coloured by medication status (yellow = schizophrenia cases not prescribed clozapine, green = treatment-resistant schizophrenia cases prescribed clozapine). The solid diagonal line depicts x=y, i.e. where the estimated and actual values are the same. The dashed diagonal line depicts the line of best fit. Presented at the top of the graph is the Pearson's correlation coefficient (r) between the estimated and actual age across all samples in that cohort. Also shown in the bottom right hand corner of each panel is an interaction P value from a test for different correlations between DNAmAge and actual age for schizophrenia patients prescribed clozapine and schizophrenia patients prescribed alternative medications.

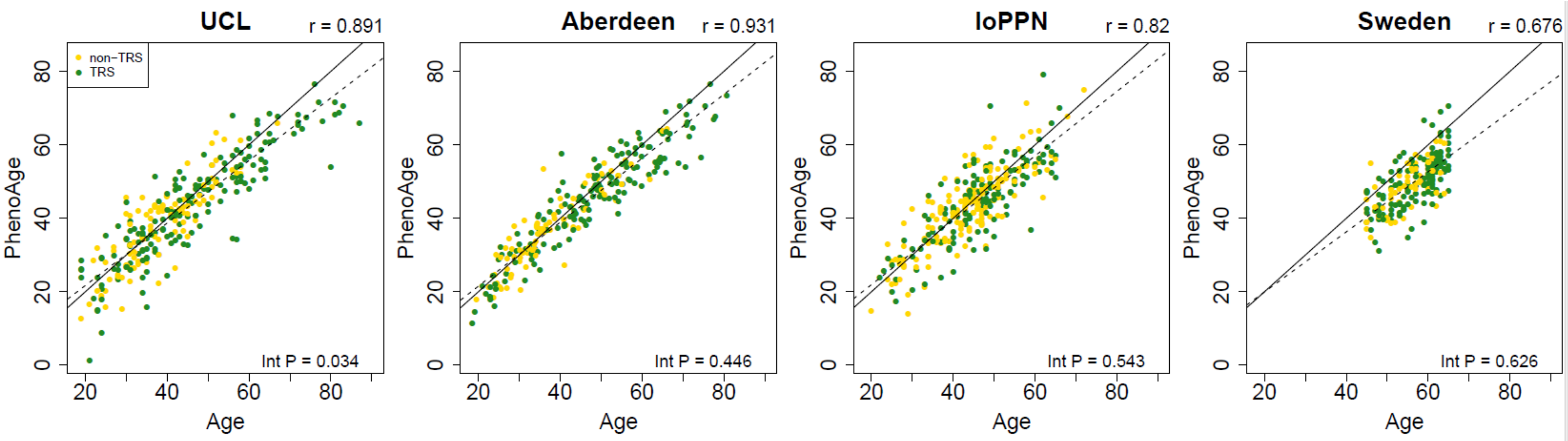


**SF10:** Scatterplots of PhenoAge derived from the DNA methylation data against actual chronological age for each of the cohorts. PhenoAge was calculated using the algorithm described by Levine et al. [1]. Each point represents an individual and is coloured by schizophrenia status (yellow = schizophrenia cases not prescribed clozapine, green = treatment-resistant schizophrenia cases prescribed clozapine). The solid diagonal line depicts x=y, i.e. where the estimated and actual values are the same. The dashed diagonal line depicts the line of best fit. Presented at the top of the graph is the Pearson's correlation coefficient (r) between the estimated and actual age across all samples in that cohort. Also shown in the bottom right hand corner of each panel is an interaction P value from a test for different correlations between PhenoAge and actual age for schizophrenia patients prescribed clozapine and schizophrenia patients prescribed alternative medications.

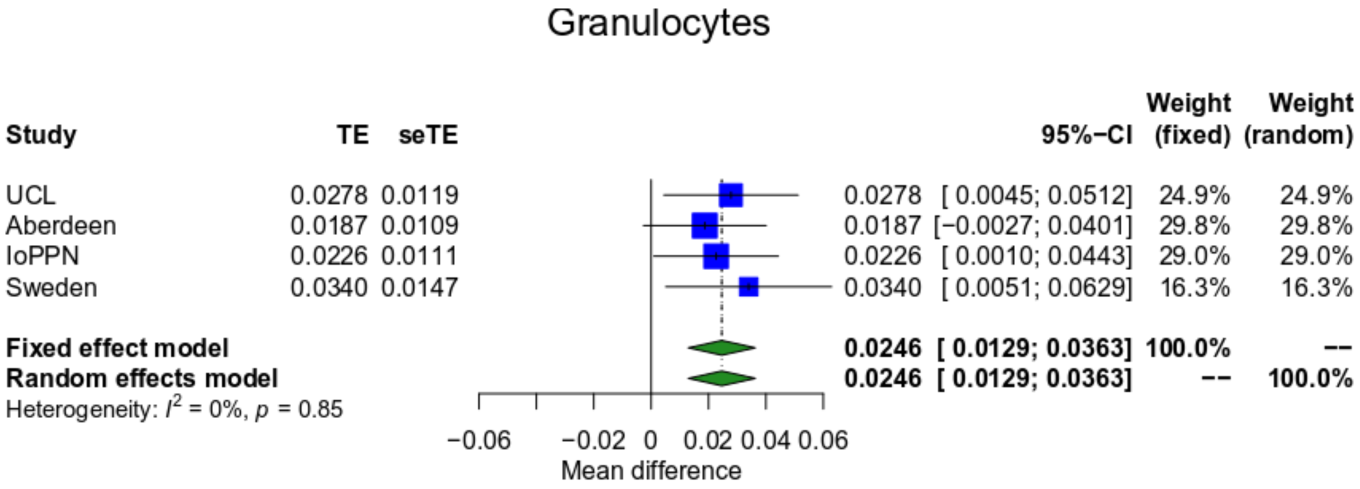


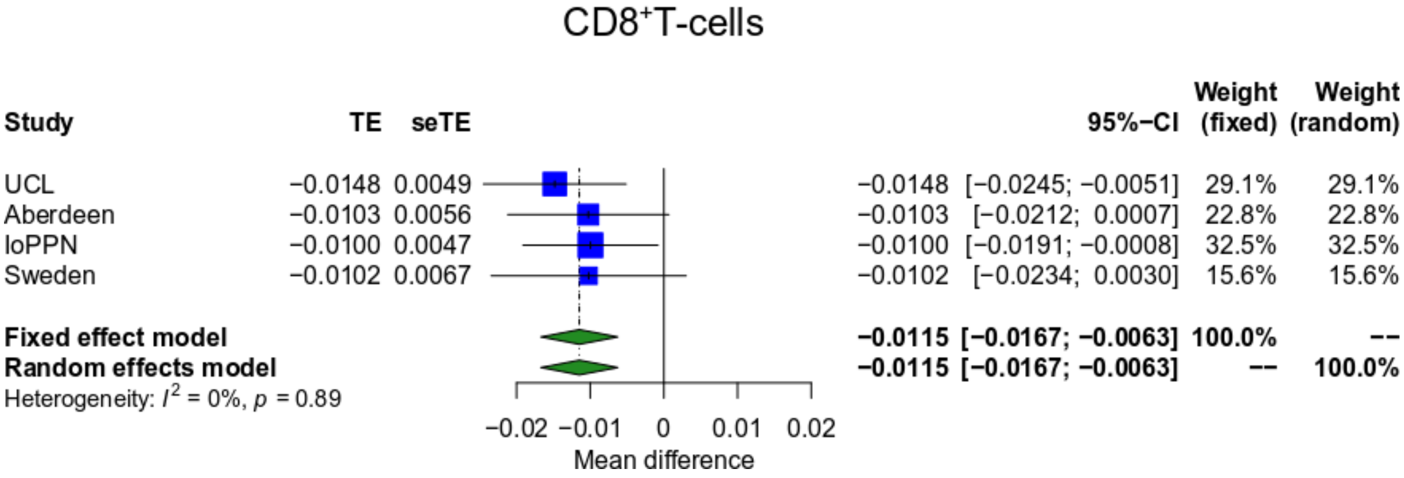


**SF11: Treatment-resistant schizophrenia patients prescribed clozapine are characterized by altered blood cell proportions.** Shown are forest plots from meta-analyses of differences in estimated blood cell proportions derived from DNA methylationdata between treatment-resistant schizophrenia patients prescribed clozapine and schizophrenia patients prescribed other medications for granulocytes, CD8+ T-cells. TE – treatment effect i.e. the mean difference between cases and controls, seTE – standard error of the treatment effect.

A

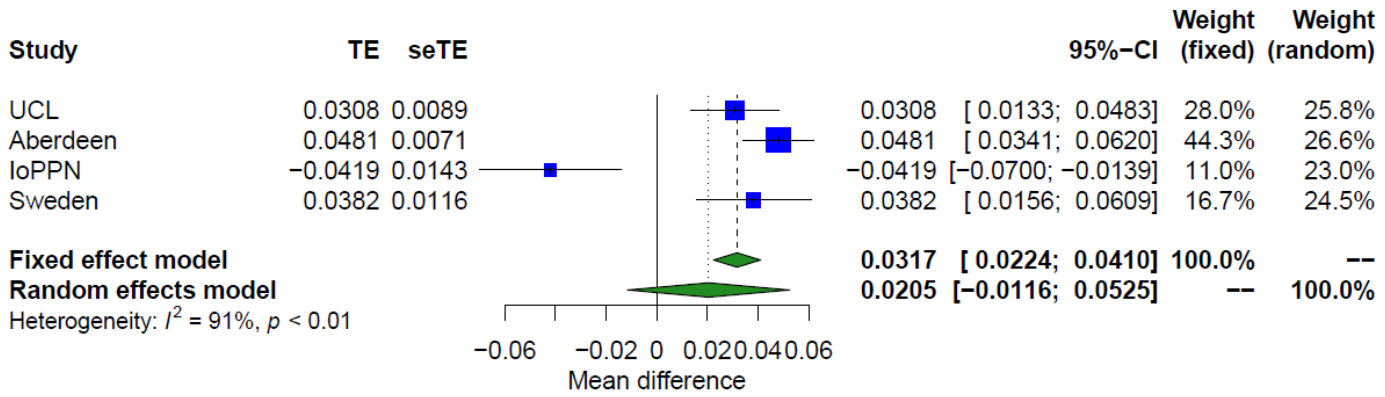


B

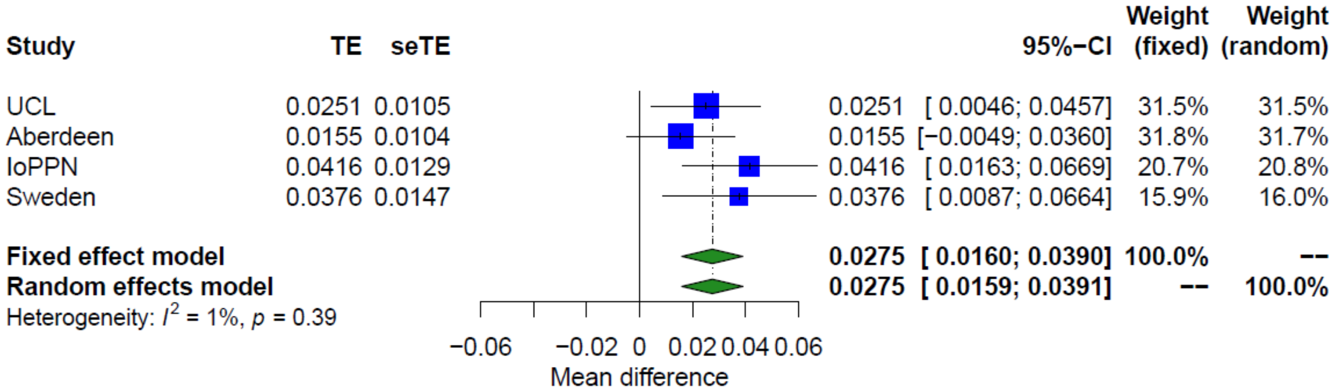


**SF12: Additive effect of schizophrenia and treatment-resistance on granulocyte proportions.** Shown are forest plots from meta-analyses of differences in estimated granulocyte proportions derived from DNA methylation data between A) schizophrenia patients and controls and B) treatment-resistant schizophrenia patients prescribed clozapine and schizophrenia patients prescribed other medications. TE – treatment effect i.e. the mean difference between cases and controls, seTE – standard error of the treatment effect.

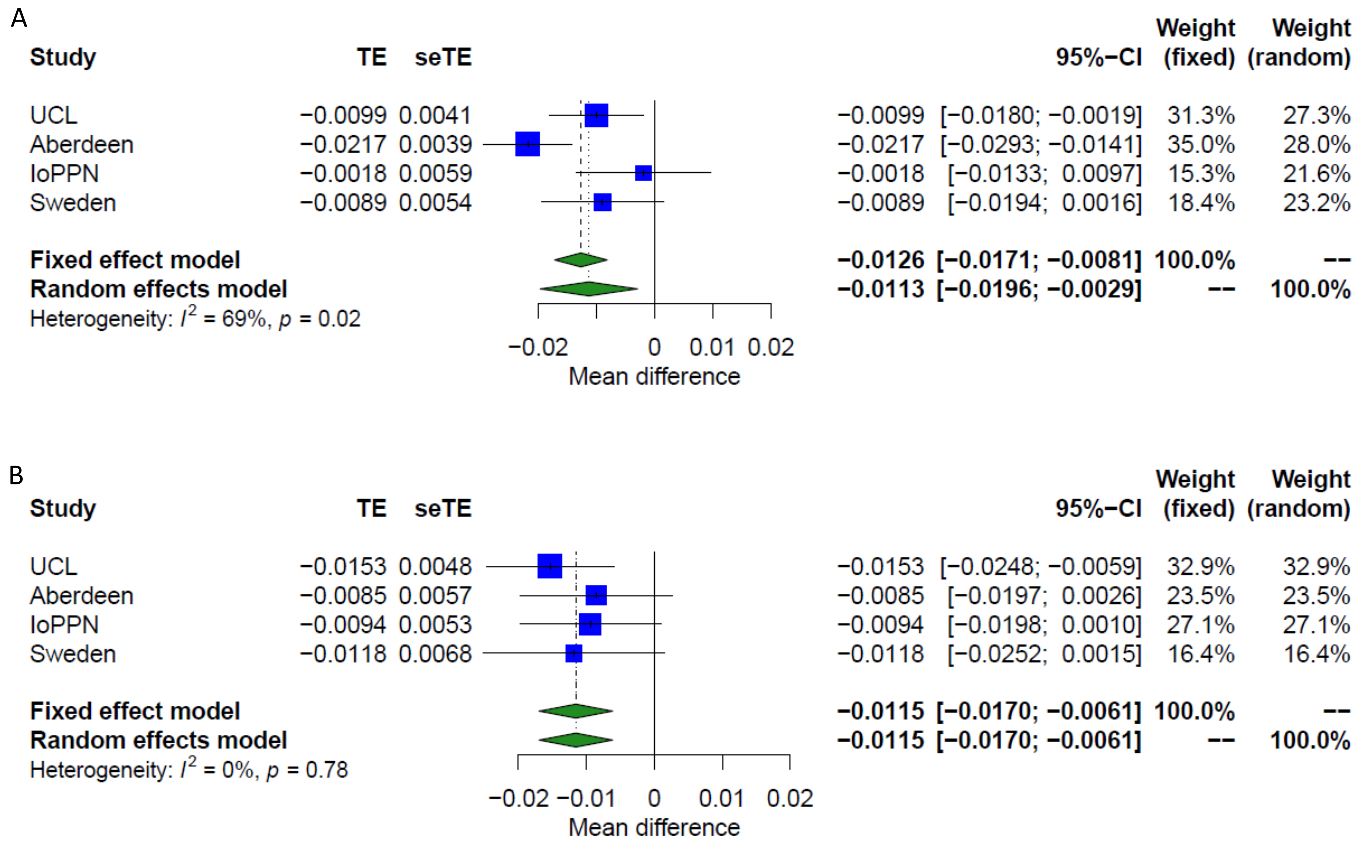


**SF13: Additive effect of schizophrenia and treatment-resistance on $CD8^+$ T-cell proportions.** Shown are forest plots from meta-analyses of differences in estimated granulocyte proportions derived from DNA methylation data between A) schizophrenia patients and controls and B) treatment-resistant schizophrenia patients prescribed clozapine and schizophrenia patients prescribed other medications. TE – treatment effect i.e. the mean difference between cases and controls, seTE – standard error of the treatment effect.

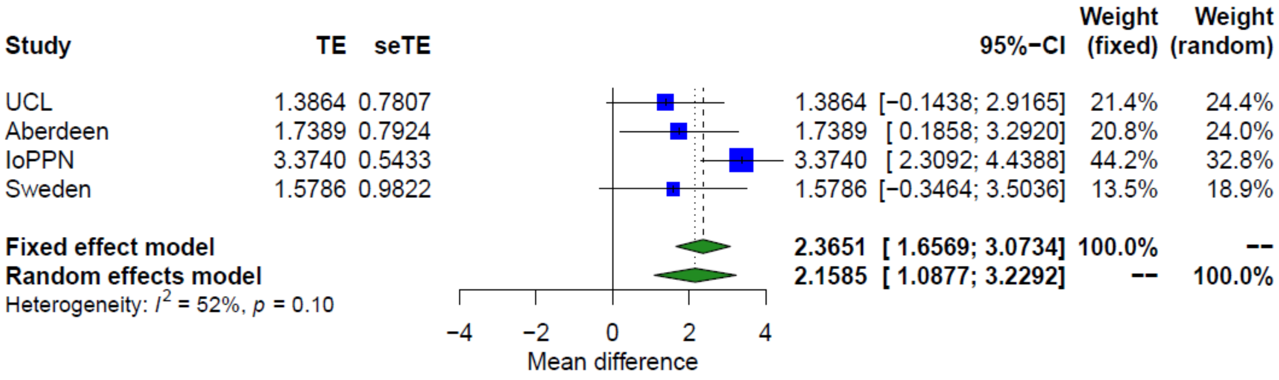


**SF14: Forest plot from meta-analyses of differences in smoking derived from DNA methylationdata between treatment-resistant schizophrenia patients prescribed clozapine and schizophrenia patients prescribed other medications.** TE – treatment effect i.e. the mean difference between cases and controls, seTE – standard error of the treatment effect.

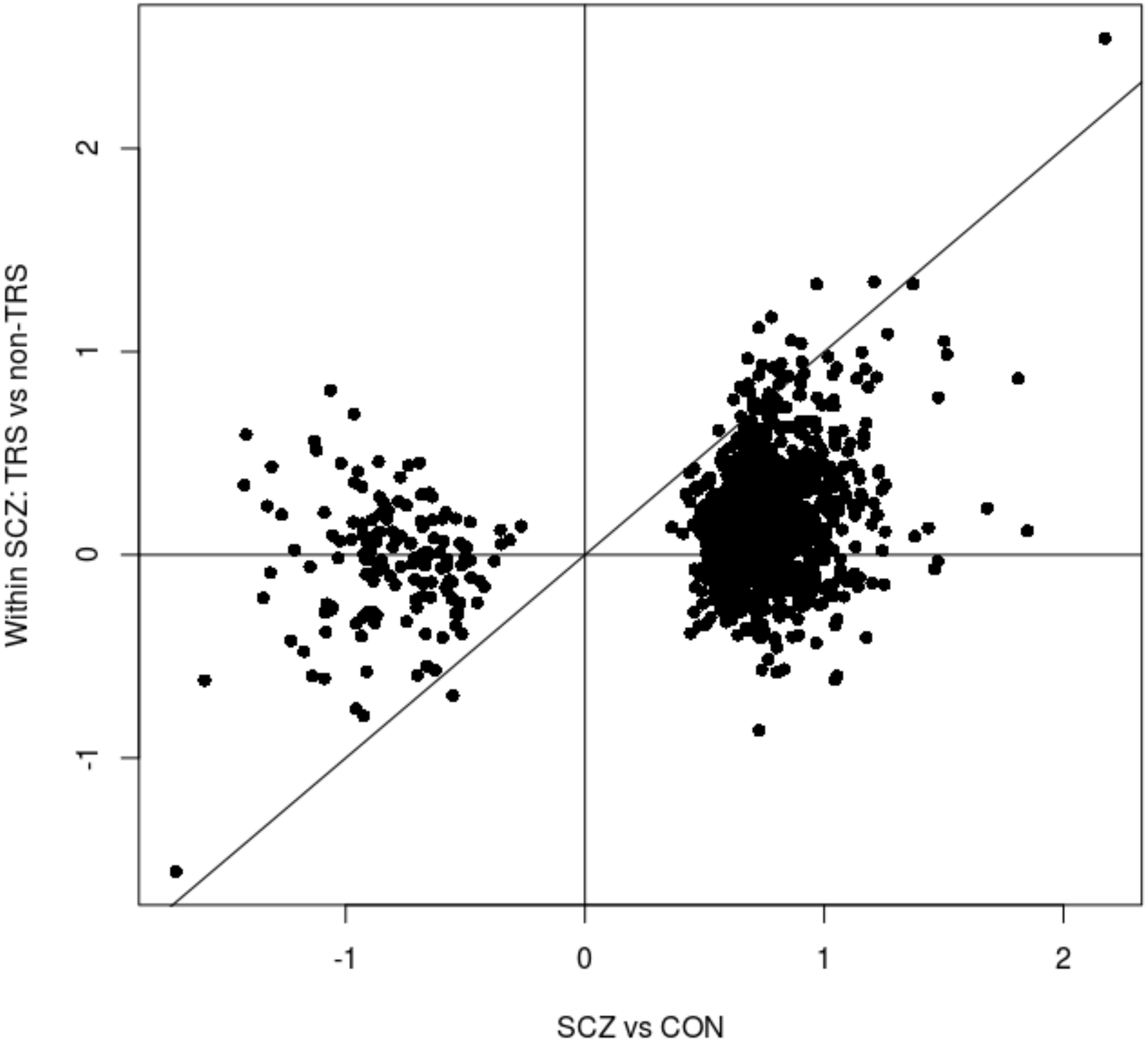


**SF15: Overlap between schizophrenia-associated DMPs and treatment-resistant-associated DMPs.** Scatterplot of schizophrenia associated DMPs (n = 1,048; $P < 9x10^{-8}$) comparing their estimated mean difference in DNA methylation between schizophrenia cases and controls (x-axis) and the estimated mean difference between treatment-resistant schizophrenia cases (TRS) and non-treatment-resistant schizophrenia cases (non-TRS) (y-axis). The solid diagonal line is x = y and indicates where the mean difference in the two analyses are equal.

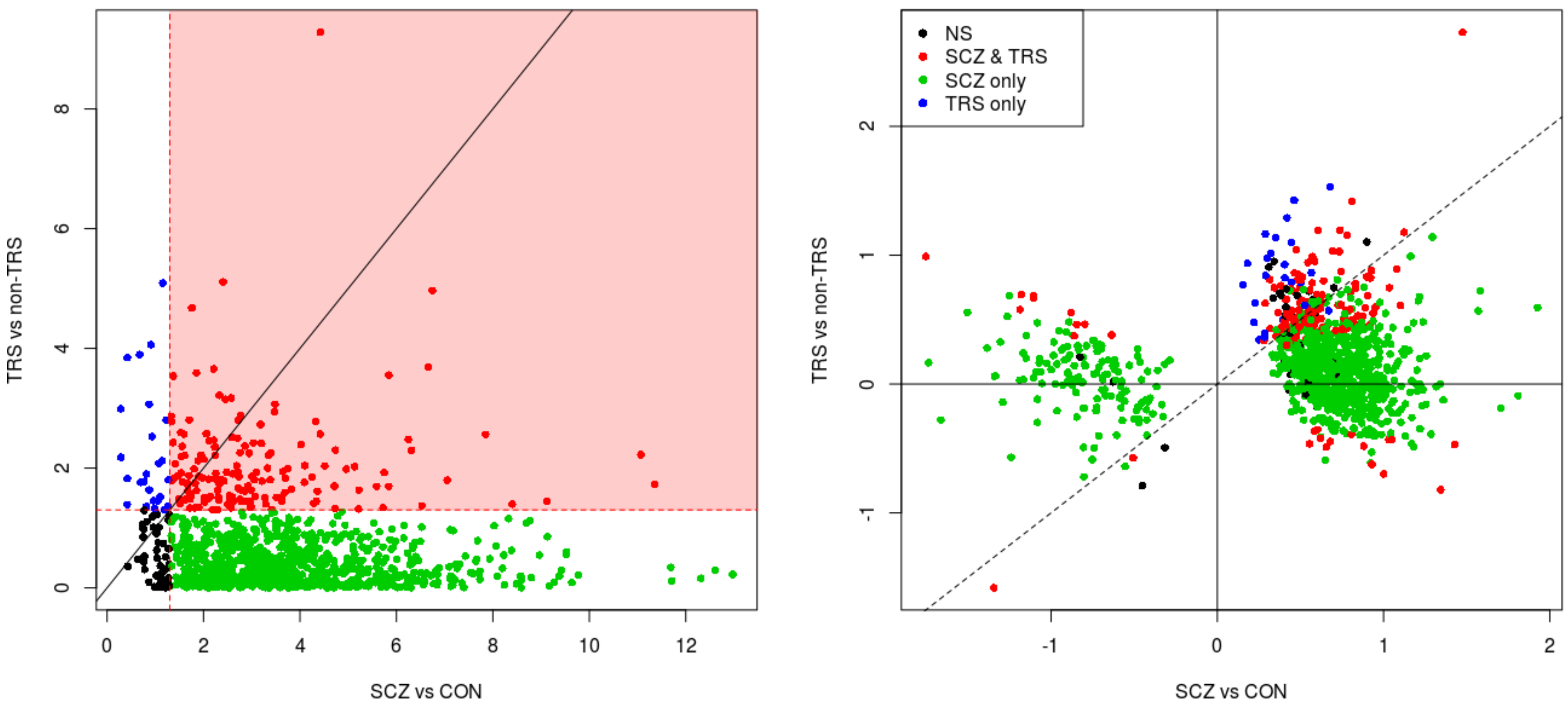


**SF16: Comparison of treatment-resistant schizophrenia effects with schizophrenia associated affects at DNA methylation sites associated with schizophrenia.** Scatterplot of the 1,048 DMPs associated with schizophrenia comparing results for schizophrenia cases and controls (x-axis) against from EWAS within schizophrenia patients comparing TRS and non-TRS. Plotted on the left is the –log10 P and on the right is the estimated mean difference, which were taken from a regression model that simultaneously quantified the effect on DNA methylation of schizophrenia status and clozapine status. The red shaded area indicates DMPs which had a significant association with clozapine (P < 0.05). NS – not significant; DMPs – differentially methylated positions; SCZ – schizophrenia; TRS = treatment-resistant schizophrenia;

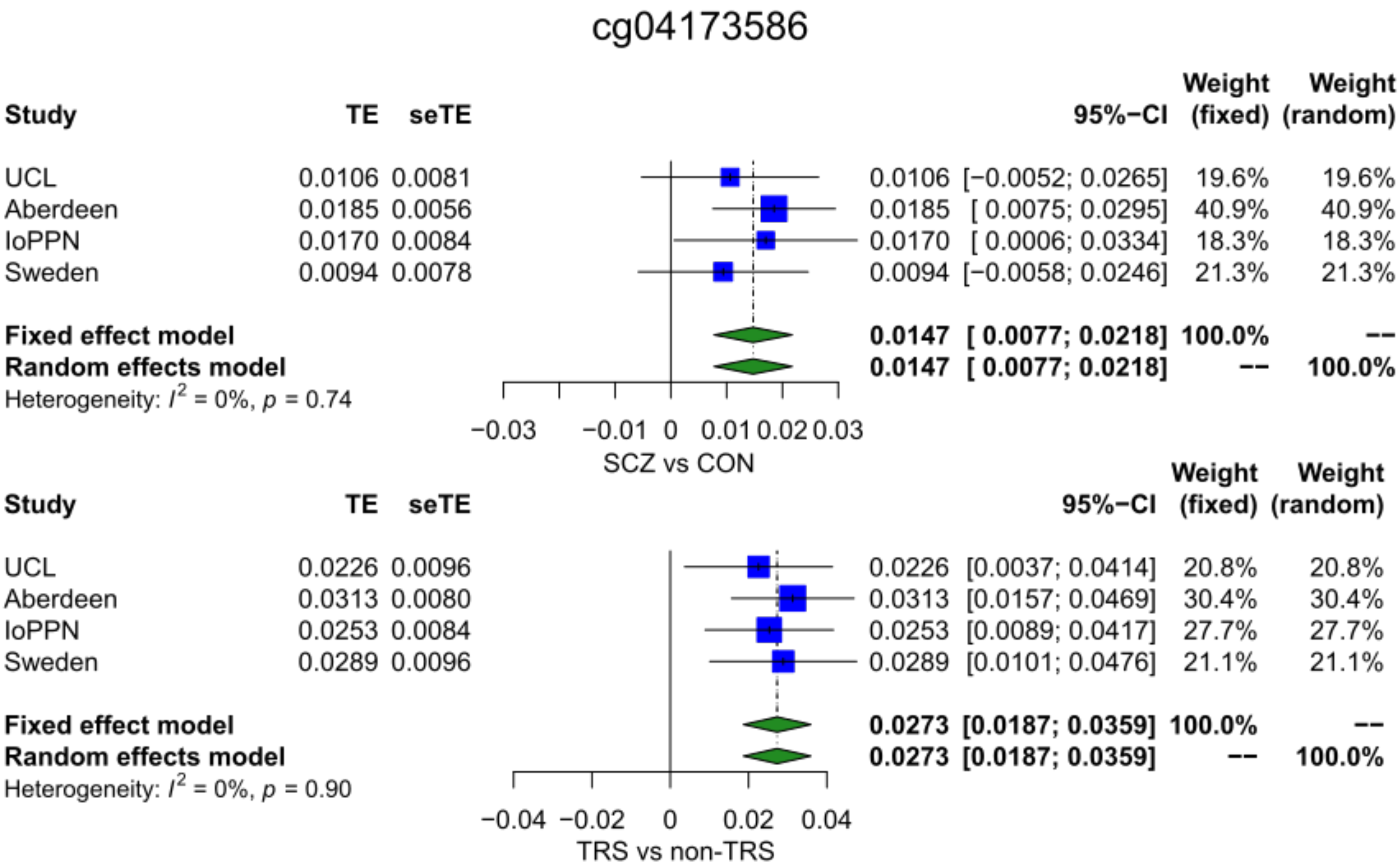


**SF17: Forest plot of a site where DNA methylation is significantly associated with schizophrenia and within cases, with treatment-resistant schizophrenia.** TE – treatment effect i.e. the mean difference between cases and controls, seTE – standard error of the treatment effect.

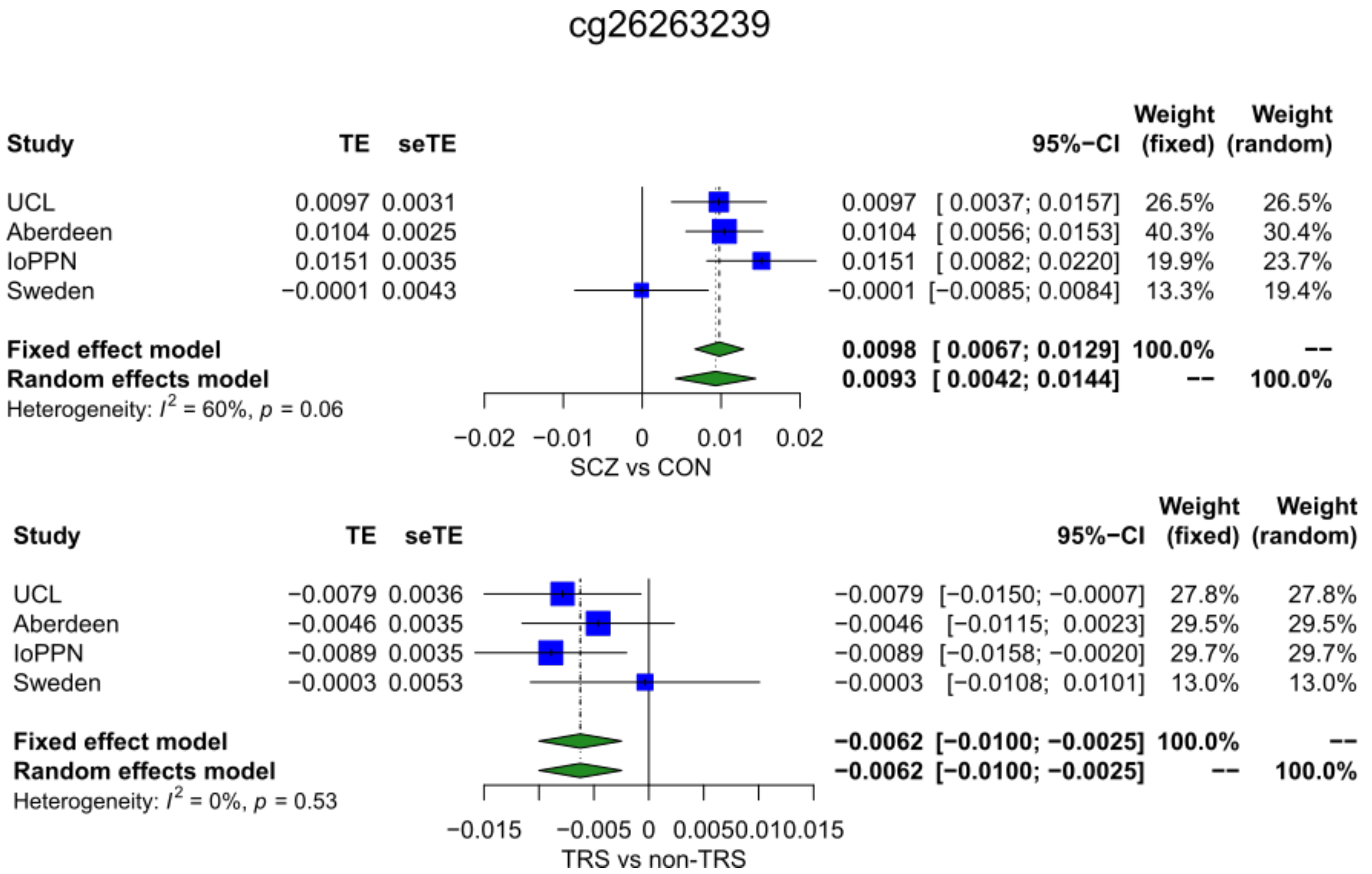


**SF18: Forest plot of a site where DNA methylation is significantly associated with schizophrenia and within cases, with treatment-resistant schizophrenia but with opposite directions of effect.** TE – treatment effect i.e. the mean difference between cases and controls, seTE – standard error of the treatment effect.
